# I-M-150847, a novel GLP-1 and GIP receptor dual agonist, improves glycemic control and reduces obesity in the rodent model of type 2 diabetes and obesity

**DOI:** 10.1101/2021.12.05.471325

**Authors:** Rathin Bauri, Shilpak Bele, Jhansi Edelli, Neelesh C. Reddy, Sreenivasulu Kurukuti, Tom Devasia, Ahamed Ibrahim, Vishal Rai, Prasenjit Mitra

## Abstract

We report the discovery of a novel unimolecular glucagon-like peptide-1 (GLP-1) and glucose-dependent insulinotropic polypeptide (GIP) receptor dual agonist that exhibits potent glycemic control and weight loss in diet-induced obese mice. The dual agonism of GLP-1 and GIP receptor was achieved by replacing the tryptophan cage of exendin-4 with the C-terminal undecapeptide sequence of oxyntomodulin along with a single amino acid substitution from histidine to tyrosine at the amino terminus of the peptide. The structural modification places lysine 30 of the novel incretin agonist in frame with the corresponding lysine residue in the native GIP sequence. The novel incretin receptor dual agonist, named I-M-150847, induces rapid redistribution of GLP-1R at the plasma membrane following activation ensuring the maintenance of the receptor in a sensitized state. I-M-150847 promotes glucose-stimulated insulin exocytosis in cultured pancreatic beta cells and augments insulin-stimulated glucose uptake in mouse adipocytes. Chronic administration of I-M-150847 enhances insulin sensitivity, improves glycemic control, and achieves significant weight loss relative to the control or exendin-4-treated DIO-mice demonstrating the therapeutic efficacy of dual agonist in ameliorating type 2 Diabetes and Obesity.

**Significance statement.:**

- Replacement of the Trp-cage with the C-terminal oxyntomodulin undecapeptide along with the tyrosine substitution at the amino terminus converts the selective GLP-1R agonist exendin-4 to a novel GLP-1R and GIPR dual agonist I-M-150847.
- The GLP-1R and GIPR dual agonist I-M-150847 induces the expeditious redistribution of GLP-1R at the plasma membrane following initial activation thereby maintaining the receptor in a sensitized state.
- The incretin receptor dual agonist I-M-150847 enhances insulin sensitivity and delivers superior glycemic control and weight loss compared to exendin-4 in the rodent model of diabetes and obesity.

**Graphical Abstract:** 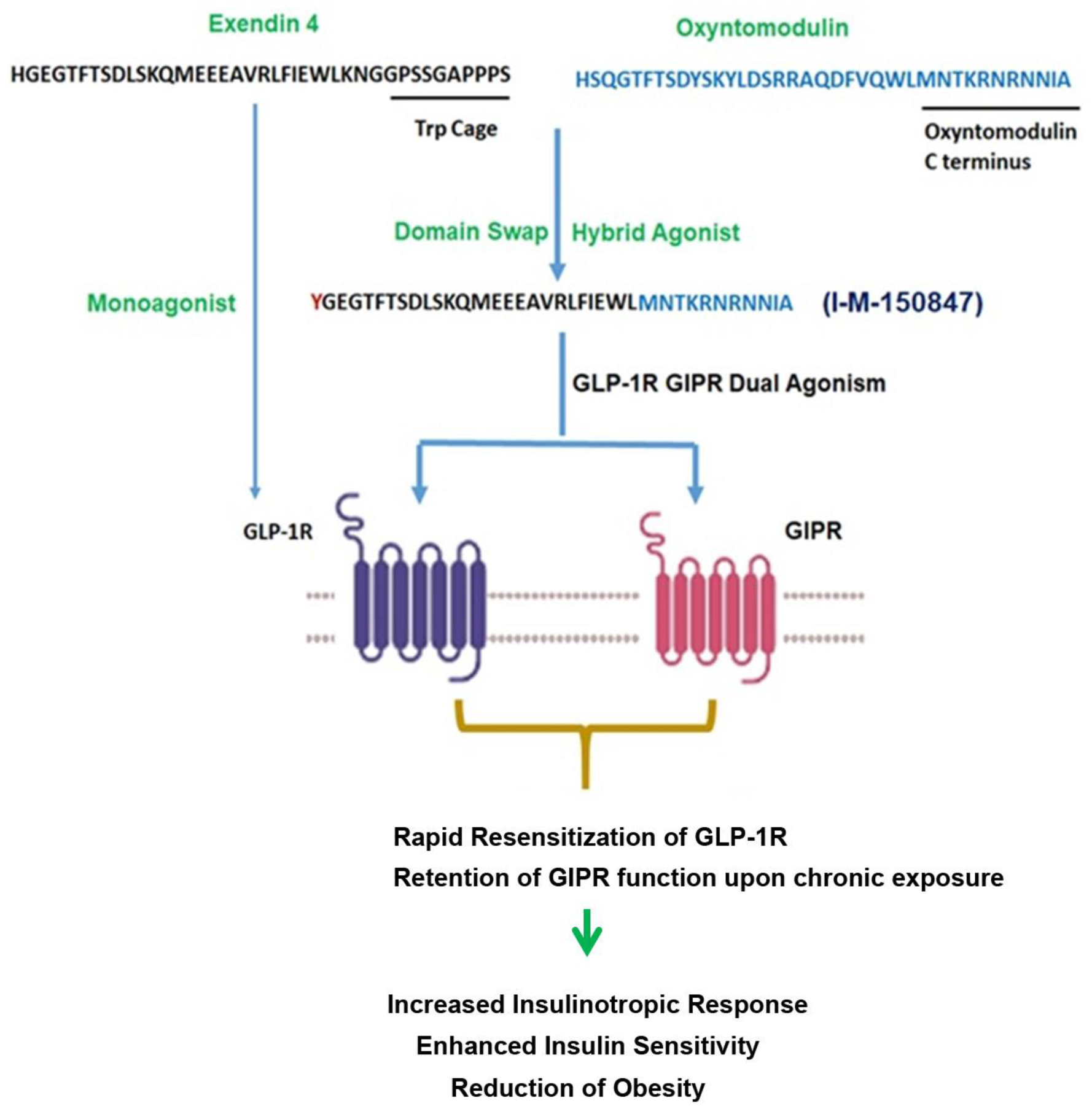

## Introduction

Incretin hormones, glucagon-like peptide-1 (GLP-1), and glucose-dependent insulinotropic polypeptide (GIP) lower the postprandial glucose as they potentiate glucose-stimulated insulin release from the pancreatic beta cells (Drucker, 2006),(Ranganath, 2008). The therapeutic efficacy of GLP-1 is presently well established and GLP-1R agonists (GLP-1RAs) have emerged as the medicine of choice for the treatment of type 2 Diabetes (T2DM) and obesity (Amori et al., 2007; Andersen et al., 2018; Astrup et al., 2009). However, not all patients receiving GLP-1RA therapy can attain long-term glycemic control and desired reduction of obesity necessitating a complementary therapeutic approach to attain proper metabolic benefits (Samms et al., 2020). The reduction of insulin resistance in the case of GLP-1RA therapy is primarily due to weight loss (Fonseca et al., 2019). In contrast, long-acting GIPR agonists (LAGIPRAs) contribute to enhanced insulin sensitivity through their action on adipose tissues (Samms et al., 2021; Varol et al., 2014). Consequently, to avail the benefits of both the incretin receptor agonists, hybrid peptides were designed incorporating residues from GLP-1 and GIP to confer dual GLP-1R and GIPR agonist activity in a single molecule (Bastin and Andreelli, 2019; DiMarchi, 2018; Finan et al., 2013). The dual agonists exhibit greater efficacy as compared to GLP-1R monoagonists in preclinical settings (Coskun et al., 2018; Finan et al., 2013) as well as in clinical trials (Frias et al., 2018),(Rizvi and Rizzo, 2022),(Jastreboff et al., 2022),(Rosenstock et al., 2021)

The approach to regulating energy balance through dual agonism exists in nature’s repository (Pocai, 2012; Pocai et al., 2009). Oxyntomodulin, an endogenous dual agonist of GLP-1R and Glucagon receptor (GCGR) displays superior bodyweight and glucose lowering effects than GLP-1R monoagonists upon enhancement of its plasma stability through chemical modifications (Kerr et al., 2010),(Muppidi et al., 2016). In a parallel approach, GIP residues were incorporated into the oxyntomodulin sequence to achieve a GIPR, GLP-1R, and GCGR triagonist response in a single hybrid molecule for the development of a potent multi-receptor agonist (Bhat et al., 2013).

In our present report, we employed a structural approach for the rational design of GLP-1R and GIPR dual agonists. The existing crystal structure reveals that the peptide ligands form alpha-helix as they bind to the extracellular domain of the incretin receptors. The affinity and specificity of the respective ligand and the receptor are determined by the interactions between the amino acid residues of the ligand and the binding site of the receptor (Parthier et al., 2007; Runge et al., 2008; Underwood et al., 2010). Our data reveal that the replacement of the tryptophan cage (trp-cage) of Exendin-4 tyrosine substituted at the amino terminus (ex-4Y1) with the C terminal undecapeptide sequence of Oxyntomodulin places the lysine 30 of the novel hybrid agonist in frame with the corresponding lysine residue in the native GIP sequence thereby converting the peptide to a GLP-1R-GIPR dual agonist. The novel incretin peptide exhibits potent GLP-1R and GIPR dual agonism with significantly diminished activity towards glucagon receptors. Sustained GLP-1R agonism from endosomes was earlier reported from our laboratory (Kuna et al., 2013), (Girada et al., 2017), In the present study, we demonstrate that I-M-150847-mediated GLP-1R signaling, contrary to exendin-4, is short-lived and facilitates rapid redistribution of GLP-1R at the plasma membrane following activation thereby maintaining the receptor in a sensitized state. Our data shows that I-M-150847 enhanced insulin sensitivity upon chronic exposure in cultured mouse adipocytes. In vivo, I-M-150847 displays superior glycemic control, enhanced insulin sensitivity, and greater weight loss compared to exendin-4 highlighting its promising efficacy for the treatment of T2DM and obesity

## Results

### Generation of potent GLP-1R and GIPR dual agonists

Our primary objective was to design a GLP-1R and GIPR dual agonist while minimizing its effect on GCGR. The crystal structure of ligand coupled to GLP-1R and GIPR extracellular domain (Parthier et al., 2007; Runge et al., 2008; Underwood et al., 2010), structure-activity relationships of incretins (Finan et al., 2013; Patterson et al., 2013; Yang et al., 2020), the similarities in their primary sequences as well as the sequence differences in the middle and the C terminal region of the respective agonists guided us to rationally design the hybrid peptide with potent GLP-1R and GIPR dual activity in a single molecule. The unimolecular GLP-1R and GIPR dual agonism was achieved in three steps. In the first step, at the C-terminus of the 9^th^ amino acid residue of I-M-150844 that corresponds to the N terminal sequence of exendin-4, we added GIP-specific amino acids Tyr^10^, Ser^11^, Ile^12^, Ala^13^, Met^14^, Asp^15^, Lys^16^, Ile^17^, His^18^, Gln^19^, Gln^20^, Asp^21^, Phe^22^, Val ^23^, Asn ^24^, Trp^25^, and Leu^26^ followed by GLP-1 specific residues Val^27^, lys^28^, Gly^29^, Arg^30^ and Gly^31^ at the C terminal end **(Table 1, Figure 1-Figure supplement 1).** The resultant peptide I-M-150845 exhibits 76.78± 6.32% GLP-1R activity, 86.86% ± 13.36 GIPR activity, and 11.5 ± 0.99 % GCGR activity measured relative to liraglutide, GIP, and oxyntomodulin in the corresponding GLP-1R. GIPR and GCGR luciferase reporter assays. In the next step, we replaced the GLP-1 specific residues Val^27^, lys^28^, Gly^29^, Arg^30,^ and Gly^31^ of I-M-150845 with the C-terminal oxyntomodulin residues Met^27^, Asn^28^, Thr^29^, Lys^30^, Arg^31^, Asn^32^, Arg^33^, Asn^34^, Asn^35^, Ile^36^, and Ala^37^ to generate a new incretin agonist I-M-150846. The peptide showed 65.65 ± 6.7 % GIPR activity but a drastically attenuated GLP-1R activity **(Table 1, Figure 1-Figure supplement 2).** To retain the GLP-1R activity, we removed the GIP-specific residues Tyr^10^, Ser^11^, Ile^12^, Ala^13^, Met^14^, Asp^15^, Lys^16^, Ile^17^, His^18^, Gln^19^, Gln^20^, Asp^21^, Phe^22^, Val ^23^, Asn ^24^ from I-M-150846 and replaced them with exendin-4 specific residues Leu^10^, Ser ^11^, Lys^12^, Gln^13^, Met^14^, Glu^15^, Glu^16^, Glu^17^, Ala^18^, Val^19^, Arg^20^, Leu^21^, Phe^22^, Ile^23,^ and Glu^24^. The novel incretin peptide I-M-150847 derived by this replacement exhibited 122.95±2.04% GLP-1R activity, 108.09 ± 5.17 % GIPR activity, and 14.93 ±1.68 % GCGR activity as described in human GLP-1R, GIPR, and GCGR cell-based cAMP reporter assay using liraglutide, GIP and Oxyntomodulin as a positive control in respective assays (**Table1, Figure 1-Figure supplement 3, Fig 1A, B, C**). To ascertain the functional characteristics of I-M-150847, we compared its potency with exendin F1 (Jones et.al 2018), and Tirzepatide, (Willard et.al 2020) in HEK 293T cell lines expressing GLP-1R. As our data shows, I-M-150847, like exendin F1 and Tirzepatide, is a full agonist of GLP-1R in cAMP accumulation assay (EC_50_ 16.35 nM) (**Fig 1D).** In HEK 293T cells expressing GIPR, I-M-150847 also displays a full agonist activity though its potency of cAMP accumulation (EC_50_ 28.90 nM) is right-shifted relative to Tirzepatide (EC_50_ 0.609 nM) **(Fig 1E).** In contrast, we observed a drastically reduced activity of I-M150847 towards GCGR in the HEK GCGR cAMP accumulation assay **(Figure1- Figure supplement 4)**. The data thus demonstrates the potent GLP-1R and GIPR dual agonism of the hybrid peptide I-M-150847 while exhibiting minimal activity towards GCGR. In cultured pancreatic beta cells, I-M-150847 shows full agonist activity and comparable potency to exendin-4 which underlines its efficacy in the physiological milieu **(Fig 1F)**. LC/MS analysis, and purity, of the C terminal amidated version of I-M-150847, are provided in **Figure1-Figure supplement 5,** and **Figure 1- Figure supplement 6.**

**Figure 1:**
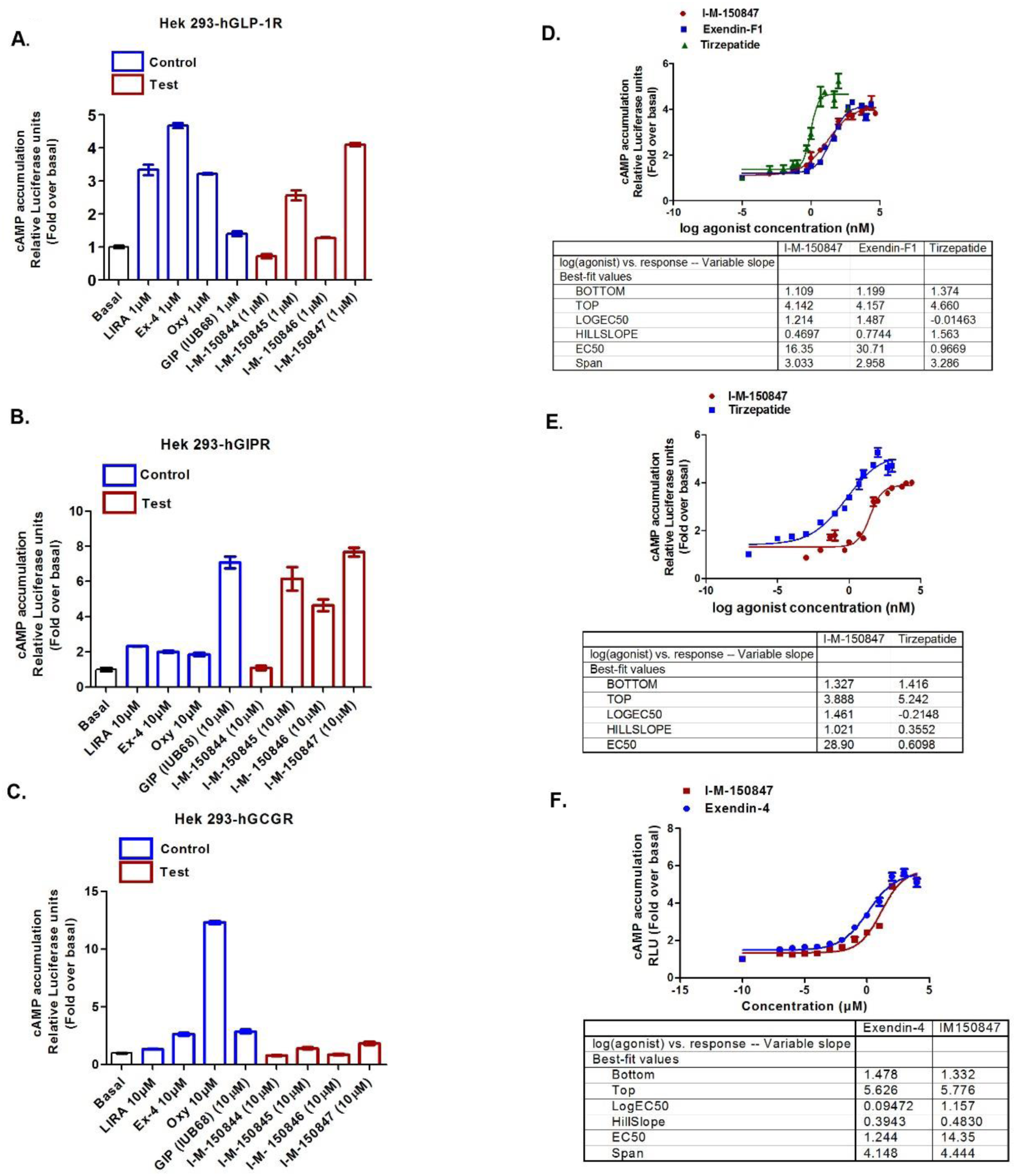
Discovery of I-M-150847 a GLP-1R GIPR dual agonist **A**. Screening of peptide analogs in HEK 293 cells expressing GLP-1R for the cAMP generation as measured by luciferase reporter assay. Y-axis represents relative Luciferase units normalized by β-galactosidase expression. The data expressed as fold over the basal is presented as means ± SD. Lira: Liraglutide; Oxm: Oxyntomodulin, EX-4: Exendin-4. **B.** Screening of peptide analogs in HEK 293 cells expressing GIPR for the cAMP generation as measured by luciferase reporter assay.**C.** Screening of peptide analogs in HEK 293 cells expressing GCGR for the cAMP generation as measured by luciferase reporter assay. **D.** Dose-response curve of stimulation of cAMP generation of I-M- 150847 vis-à-vis Exendin-F1 and Tirzepatide in HEK 293 cells expressing GLP-1R as measured by cAMP responsive element luciferase reporter assay. EC_50_ represented as mean ±SE of three independent experiments.**E.** Dose-response curve of stimulation of cAMP generation of I-M-150847 vis-à-vis Tirzepatide in HEK 293 cells expressing GIPRR as measured by cAMP responsive element luciferase reporter assay. EC_50_ is represented as the mean ±SE of two independent experiments carried out in duplicate. **F.** Dose-response curve of stimulation of cAMP generation of I-M-150847 relative to exendin-4 in BRIN-BD11 pancreatic beta cells as measured by cAMP responsive element luciferase reporter assay. EC_50_ is represented as the mean ±SEM of three independent experiments carried out in duplicate.

**Table 1:**
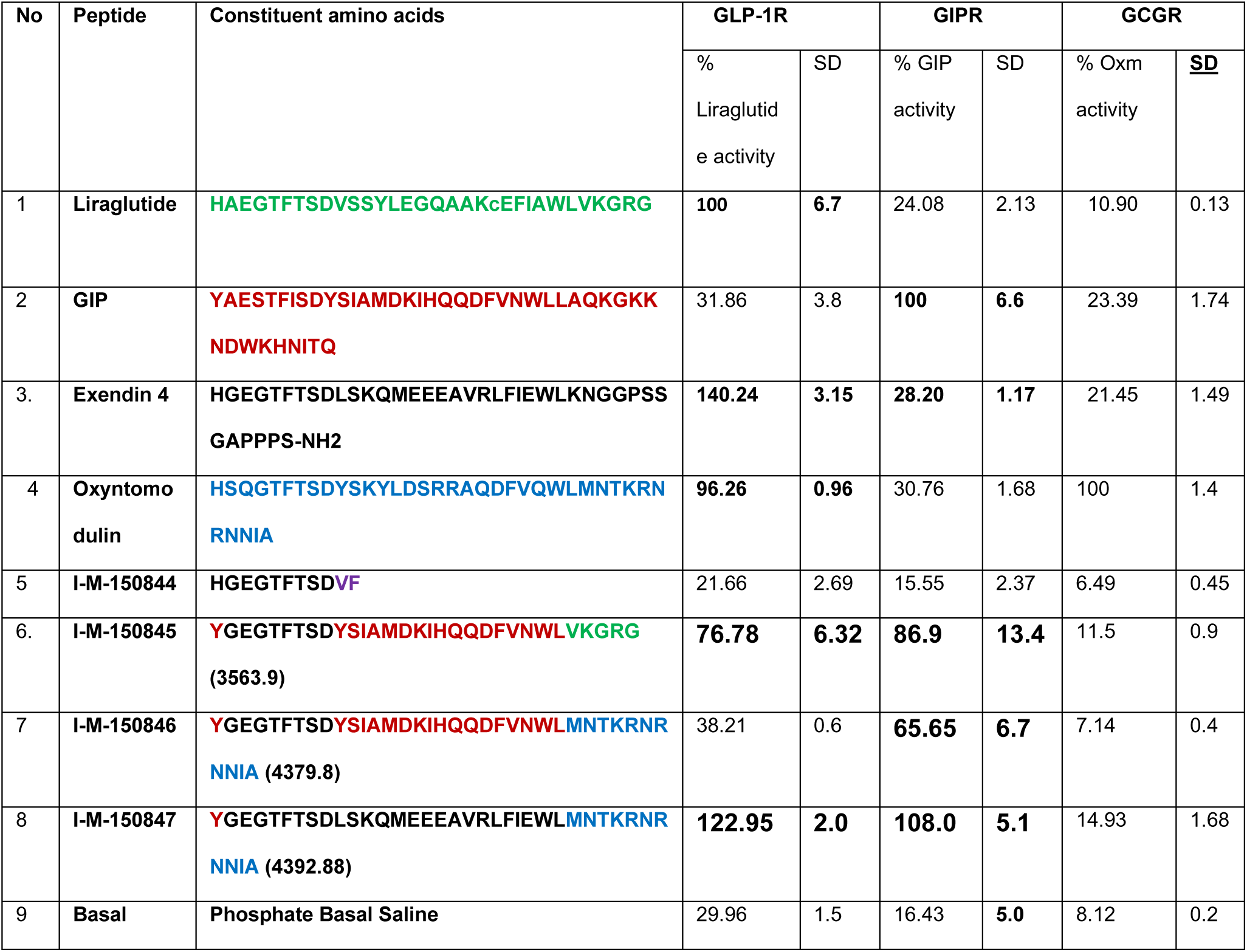
In vitro GLP-1R, GIPR, and GCGR activity of peptide analogs. The activity of the peptide analogs in vitro in GLP-1R, GIPR, and GCGR assay system as measured by cyclic AMP responsive element luciferase reporter assay. % Liraglutide activity is defined as the normalized relative luciferase units generated upon treatment of HEK –GLP-1R expressing cells with 1µM liraglutide. % GIP activity is defined as the normalized relative luciferase units generated upon treatment of HEK –GIPR expressing cells with 10µM GIP. % GCGR activity is defined as the normalized relative luciferase units generated upon treatment of HEK –GCGR expressing cells with 10µM Oxyntomodulin. Oxm =Oxyntomodulin. The numbers in brackets represent the molecular mass of the peptide. The data represent the means ± SD of the experiment carried out in duplicate in each heterologous assay system. The amino acids that correspond to Liraglutide, exendin-4, oxyntomodulin, and GIP are colored green, black, blue, and red respectively.

### I-M-150847 elicits short-term GLP-1R-mediated cAMP generation and facilitates expeditious redistribution of the functional receptor at the plasma membrane

GLP-1RAs exhibit variable efficiencies of internalization and trafficking (Jones et al., 2018). We first reported the prolonged internalization of GLP-1R upon activation that is associated with sustained GLP-1R mediated cAMP signaling upon treatment with GLP-1-conjugated to tetramethyl rhodamine (GLP-1Tmr), liraglutide, and exendin-4 (Kuna et al., 2013), (Girada et al., 2017). In our present study, we compared the trafficking and signaling efficacy of I-M-150847 relative to GLP-1R monoagonist exendin-4 in cultured pancreatic beta cells. We observed that, unlike exendin-4, cAMP generation upon treatment with I-M-150847 is short-lived, as it maximizes at 10 minutes of activation and recedes abruptly at 30 min after the agonist treatment **(Fig 2A).** Along with the short-term cAMP generation, our data reveals contrasting post-endocytic sorting of GLP-1R upon I-M-150847 treatment relative to exendin-4 **(Fig 2B).** At 0 min, we observed an annular ring of GLP-1R GFP at the plasma membrane and a diffused distribution at the cytoplasm **(Fig 2B i).** At 10 minutes following the treatment with exendin-4, there is a complete loss of the annular ring as the GLP-1R GFP signal appears as punctate dots in the cytoplasm demonstrating complete internalization of the receptor after activation with the agonist. In contrast, when treated with I-M-150847 for 10 minutes, we observed the presence of GLP-1R -GFP both as an annular ring at the plasma membrane and as punctate dots in the cytoplasm (**Fig 2B ii)**. Following 30-minute treatment with exendin- 4, the punctate dots representing GLP-1R GFP is prevalent in the cytoplasm whereas in I-M150847 treated cells, we observe the prominent annular ring at the plasma membrane as well as a few GLP-1R GFP punctate dots at the cytoplasm mostly distributed at the periphery near the plasma membrane **(Fig 2B iii).** The punctate GLP-1R GFP dots are retained in the cytoplasm after 60 minutes of exendin-4 treatment; however, in I-M-150847 treated cells, we observe the majority of GLP-1R -GFP fluorescence at the plasma membrane as annular ring; the punctate GLP-1R GFP dots are progressively much less in number with time and have a limited distribution in the proximity of the plasma membrane **(Fig 2B iv).** The data thus demonstrates that I-M-150847 exhibits reduced internalization and /or faster recycling of GLP-1R compared to exendin-4 as the maximum GLP-1R-GFP signal is observed back at the plasma membrane on a much shorter time course following activation with I-M-150847.

**Figure 2:**
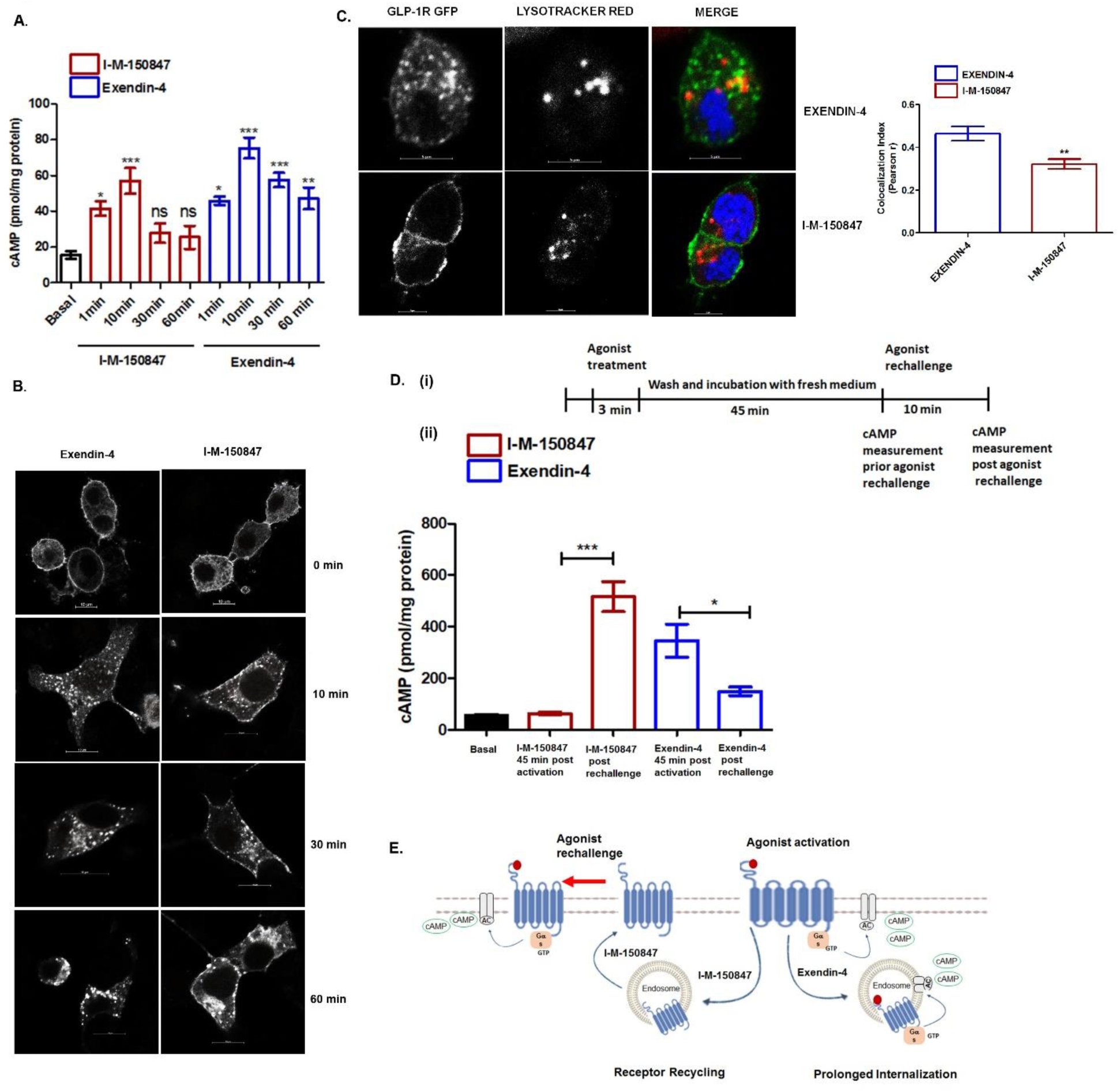
I-M-150847 facilitates prompt redistribution of GLP-1R at the plasma membrane following activation. **A.** The time-course of cAMP accumulation in BRIN-BD11 pancreatic beta cells following activation by I-M-150847 or exendin-4. The cells were incubated with 1µM I-M-150847 or exendin-4 and 3 minutes after the incubation the excess ligand was washed with KRB buffer. The cAMP was measured at 1 min, 10, 30, and 60 min after KRB wash using Direct cAMP Enzyme Immunoassay. The data are expressed as mean ±SE of three independent experiments *p<0.05; ***p<0.001, one way-ANOVA Tukey’s multiple comparison test, comparing cAMP generation upon agonist treatment relative to basal cAMP accumulation. **B.** Confocal images studying GLP-1R internalization at different time points upon exendin-4 or I-M-150847 treatment, representative image from 4 independent experiments, n=15 cells each time point, scale bars 10µM. **C**. Colocalization of GLP-1R -GFP with lysotracker red 45 min after activation with exendin-4 or I-M-150847 as evidenced in confocal microscopy, representative image from 2 independent experiments, n=5 for each experiment. Image quantification was carried out in Image J (JACoP plugin) and co-localization was measured by Pearson correlation coefficient (r) represented as mean ± SE; **p<0.01, n=5, student’s t-test (unpaired) comparing correlation coefficient (Pearson r) between exendin-4 and I-M-150847 treated cells. **D;(i)** Schematic diagram of the agonist rechallenge in BRIN-BD11 pancreatic beta cells. The cells were subjected to treatment with exendin-4 or I-M-150847 for 3 minutes following which the cells were washed and incubated for another 45-minute following which they are rechallenged with exendin-4 or I-M-150847 and cAMP generation was measured; **(ii)** Agonist rechallenge experiment; the y-axis represents cAMP accumulation (pmol/mg protein), the data is expressed as mean ± sem; *p<0.05, ***p<0.001, one-way ANOVA, n=3 independent experiments, Tukey’s multiple comparison test comparing cAMP accumulation by agonists pre and post rechallenge. **E:** Schematic illustration of GLP-1R trafficking upon activation by exendin-4 or I-M-150847.

We first reported lysosome as one of the destinations of GLP-1R following internalization of the receptor (Kuna et al., 2013) which was supported by subsequent studies using immunoelectron microscopy (Jones et al., 2018). However endocytic destination of GLP-1R was later found to be ligand specific as the receptor has been shown to recycle back to the plasma membrane upon treatment with Exendin F1 (Jones et al., 2018). In our present study, we compared the association of GLP-1R with lysosomes at 45 minutes post-treatment with exendin-4 or I-M-150847. We observed partial co-localization of GLP-1R -GFP with lysotracker red (colocalization index, Pearson r=0.46 ±0.03) upon treatment with exendin-4 at 45 minutes post internalization of the receptor **(Fig 2C).** In contrast, the localization of GLP-1R to lysosome was significantly reduced upon treatment with I-M-150847 (colocalization index, Pearson r=0.32 ±0.23; **p<0.01, n=5) **(Fig 2C).** To functionally assess the GLP-1R trafficking we next carried out the receptor resensitization experiment **(Fig 2D).** The pancreatic beta cells were treated with exendin-4 or I-M-150847 for 3 minutes followed by a wash with KRB buffer. After 45 minutes of treatment, the cells were rechallenged with respective agonists for 10 minutes and cAMP accumulation was evaluated. As **Fig 2D** shows, cultured pancreatic beta cells with prior exposure to I-M-150847 respond more efficiently to GLP-1R mediated cAMP generation relative to the cells that are pre-exposed to exendin-4 during agonist rechallenge. We reported that I-M-150847-mediated cAMP generation subsided 30 minutes post-treatment **(Fig 2A).** Aligned to these findings, cAMP generation is attenuated to the basal level 45 minutes after treatment with I-M-150847. Thereafter, upon the I-M-150847 rechallenge, we observed an increase in cAMP generation (from 61.30 ± 11.96 to 342.3 ±61.68 pmol/mg protein, 5.58-fold increase, **p<0.01, one-way ANOVA, Tukey’s multiple comparison test, n=3). The increase of cAMP generation upon rechallenge signifies prompt redistribution of the functional GLP-1R at the plasma membrane after initial activation. In contrast, similar exendin-4 treatment resulted in a sustained cAMP generation that decreases from 255.3± 54.91 to 143.8 ± 33.39 pmol/mg protein, (n=3) after agonist rechallenge. The data taken together demonstrate that the I-M-150847 treatment ensures efficient receptor recycling that preserves a higher level of functional GLP-1R at the plasma membrane relative to exendin-4 sufficient to initiate de novo second messenger signaling upon I-M-150847 re-exposure. **Fig 2E** explains the mechanism of GLP-1R trafficking upon activation by I-M-150847 vis-à-vis exendin-4.

### Prolonged I-M-150847 exposure has a contrasting functional response in cultured pancreatic beta cells and mouse adipocytes

We next evaluated whether the rapid redistribution at the plasma membrane has any functional advantage over prolonged internalization of GLP-1R following activation by agonists upon continuous exposure. As **Fig 3A** reveals, I-M-150847 significantly enhances GSIS relative to exendin-4 after 30 minutes of incubation (33.01±1.94 fold over basal upon I-M-150847 compared to 24.04 ±0.71 fold over basal upon exendin-4 treatment, **p<0.01, n=3) demonstrating its superior insulinotropic property. However, upon prolonged exposure, we observed blunting of GSIS both upon exendin-4 and I-M-150847 treatment **(Fig 3A).** The data thus underlines that GLP-1R induced GSIS in BRIN-BD11 pancreatic beta cells is independent of the GLP-1R sorting dynamics as both prolonged internalizations upon exendin-4-mediated activation or rapid redistribution at the plasma membrane following I-M-150847 treatment has a similar impact on GLP-1R desensitization upon chronic agonist exposure. The GSIS response in BRIN-BD11 pancreatic beta cells is mostly GLP-1R specific; as we observed a 6.95-fold enrichment of GLP-1R expression over GIPR in BRIN-BD11 pancreatic beta cells **(Figure 3- Figure supplement 1)** In contrast, in cultured mouse adipocytes where GIPR is predominant (McIntosh et al., 2012), the I-M-150847 treatment increases the cAMP generation from 31.6±3.7 pmol to 72.44±2.535 pmol per mg protein, (***p< 0.001, one-way ANOVA Tukey’s multiple comparison test n=3) upon chronic exposure **(Fig 3B)**. Since enhanced cAMP generation is indicative of increased lipolysis and subsequent blunting of insulin sensitivity, we assessed insulin-stimulated glucose uptake in 3T3L1 adipocytes following 16 h treatment with exendin-4 or I-M-150847. Contrary to our postulation, we observed an increase in insulin-stimulated glucose uptake upon I-M-150847 treatment (from 1.385±0.009 fold to 1.91±0.05 fold, ***p<0.001, one-way ANOVA, Tukey’s multi comparison test n=3) **(Fig 3C)** thereby demonstrating the efficacy of the incretin receptor dual agonist I-M-150847 in enhancing insulin sensitivity in cultured adipocytes. The data thus signifies that the functional outcome of chronic exposure to I-M-150847 is different in mouse adipocytes and cultured pancreatic beta cells. The data further demonstrates that the enhancement of insulin sensitivity upon I-M-150847 treatment is independent of the agonist-induced cAMP response in cultured mouse adipocytes.

**Figure 3:**
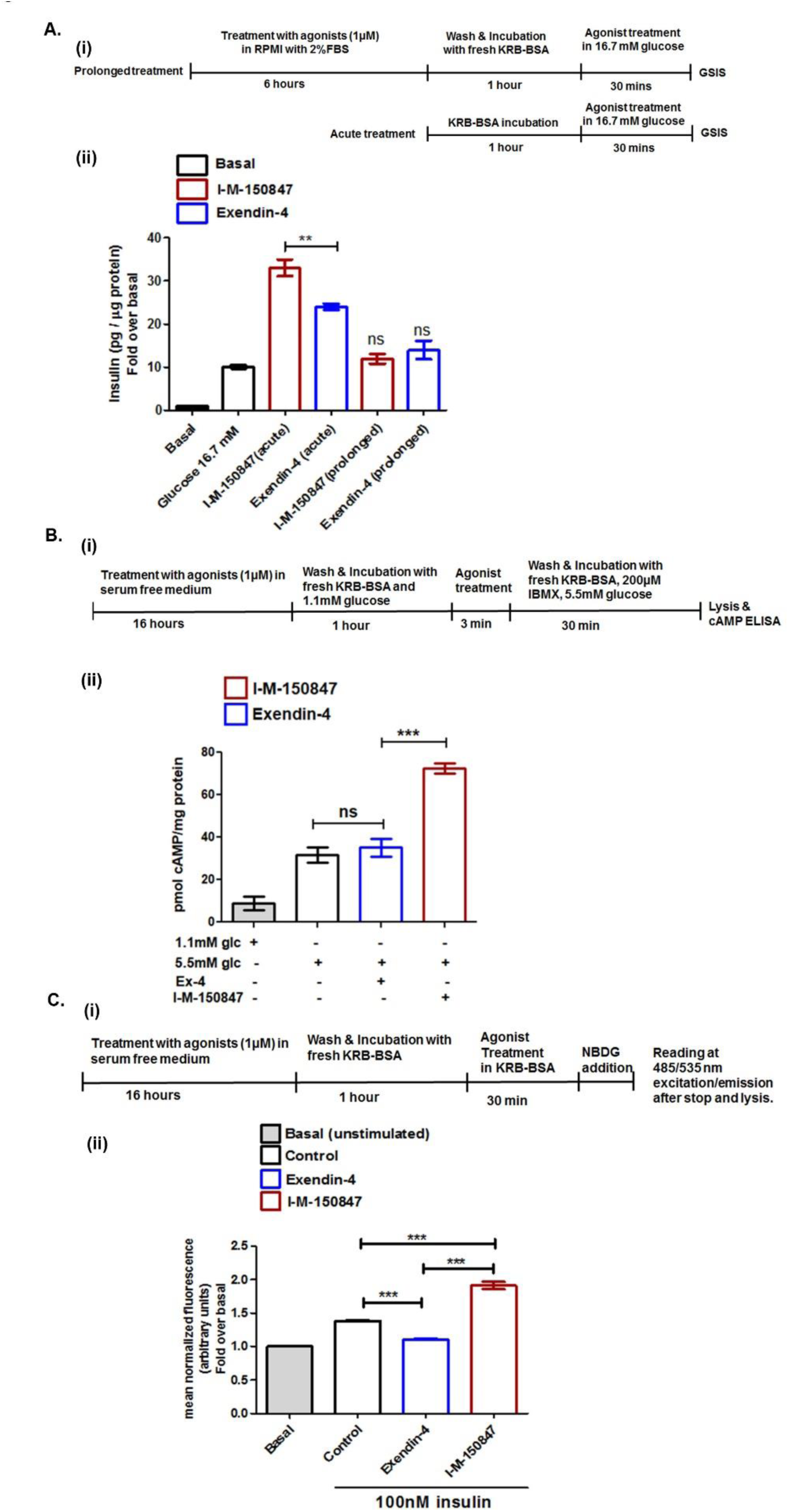
Functional response of I-M-150847 in cultured pancreatic beta cells and mouse adipocytes. **A. (i)** Schematic of GSIS in BRIN-BD11 pancreatic beta cells upon acute and chronic treatment with exendin-4 or I-M-150847. **(ii)** Acute and prolonged GSIS upon exendin-4 or I-M-150847 treatment. Y axis represents insulin secretion (pg./mg protein) expressed as the fold increase over basal insulin secretion at 0.1mM glucose. The data are expressed as mean ± SE **p<0.01, one-way ANOVA, n=3 Tukey’s multiple comparison test comparing GSIS upon 30 min treatment (acute mode) with exendin-4 and I-M-150847; ns=non-significant comparing GSIS upon 6h (prolonged) exendin-4 or I-M-150847 treatment with insulin secretion at 16.7mM glucose. **B (i)** Schematic of I-M-150847 or exendin-4 mediated cAMP generation upon overnight pre-exposure with agonists. **(ii)** The cAMP generation upon exendin-4 or I-M- 150847 treatment following overnight pre-exposure with agonists. The y axis represents cAMP accumulation (pmol/mg protein), the data are expressed as mean ± SE; ***p<0.001, one-way ANOVA, n=3, Tukey’s multiple comparison test comparing cAMP generation between exendin-4 and I-M-150847 treatment, ns=non-significant comparing prolonged cAMP generation upon agonist treatment with untreated control cells. **C(i)** Schematic of insulin-stimulated glucose uptake upon overnight pre-exposure with exendin-4 or I-M-150847 in 3T3-L1 adipocytes; **(ii)** Insulin-stimulated glucose uptake upon overnight pre-exposure with exendin-4 or I-M-150847 in 3T3-L1 adipocytes; the data are expressed as mean ± SE; ***p<0.001, n=3, one-way ANOVA, Tukey’s multiple comparison test.

### I-M-150847 outperforms Exendin-4 in imparting glycemic benefits and reducing obesity

To explore whether these in vitro observations translate to better regulation of glucose homeostasis and weight loss in the rodent model of diabesity, we compared the effect of I-M-150847 and exendin-4 in regulating hyperglycemia and obesity in C57BL/6 mice fed on the high-fat diet. We carried out twice-daily dosing of 100nmol/ kg body weight of exendin-4 or I-M-150847 for 28 days and assessed the metabolic parameters as described in the flow diagram **(Fig 4A).** As our data reveals, 10 days of exendin-4 treatment reduces fasting blood glucose from 176.5 ±11.31 mg/dL glucose to 130 ±3.02 mg/dL compared to vehicle control (***p<0.001, one-way ANOVA, Tukey’s multi comparison test n=6) **(Fig 4B (ii)).** In comparison, after 10 days of I-M-150847 treatment, fasting blood glucose was reduced from 183 ±14.02 mg/dL glucose to 101.7 ±2.17 mg/dL glucose (***p<0.001, one-way ANOVA, Tukey’s multi comparison test, n=6) **(Fig 4B (iii**)). The results thus reveal that after 10 days of treatment, I-M-150847 outperforms exendin-4 in reducing fasting blood sugar in DIO mice (mean blood glucose 101.7±2.17 mg/dL in I-M-150847 treated group versus 130 ±3.02 mg/dL exendin-4 treated group, ***p<0.001, one-way ANOVA, Tukey’s multi comparison test, n=6 **(Fig 4B (iv))**. A comparative glucose tolerance test on the 20^th^ day of agonist treatment also demonstrates the superiority of I-M-150847 over exendin-4 in executing glycemic control. As **Fig 4C** reveals, glucose excursion is superior upon I-M-150847 treatment relative to exendin-4 mediated therapy (mean AUC 20809 ± 312.1, in exendin-4 treated group, versus mean AUC 13218 ± 164.5, in I-M-150847 treated group; p<0.001, one-way ANOVA, Tukey’s multi comparison test, n=6). To explore whether enhanced glucose disposal is associated with increased insulin sensitivity, we carried out the insulin tolerance test. I-M-150847 treatment, in our study, displayed better insulin sensitivity compared to exendin-4 treatment (AUC 6149 ± 125.7 upon exendin-4 treatment, versus AUC 5134 ± 39.54 upon I-M-150847 treatment; ***p<0.001, one-way ANOVA, Tukey’s multi comparison test, n=6) as demonstrated in **Fig 4D**. The data, taken together, underlines the superiority of the dual agonist I-M-150847 over exendin-4 mediated therapy in regulating glucose homeostasis.

**Figure 4:**
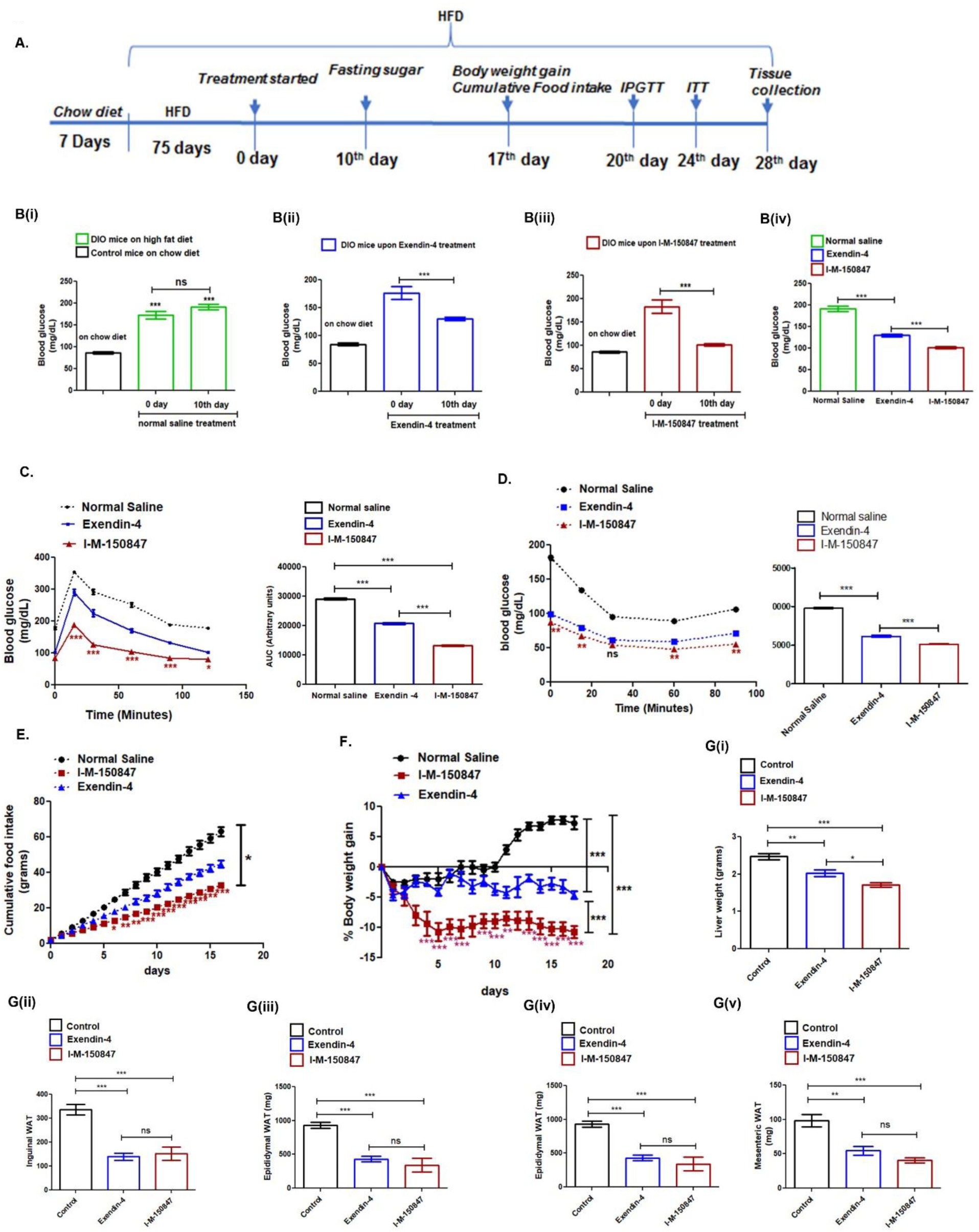
Comparison of I-M-150847 and exendin-4 in providing glycemic benefits and reducing obesity in DIO mice. **A.** Schematic diagram of the diet and the treatment paradigm as well as the pharmacological parameters measured upon the treatment of I- M-150847 and exendin-4. **B.** Measurement of the fasting blood glucose at 0 day versus the 10^th^ day of treatment; **(i)** Fasting blood glucose in control mice on high fat diet at 0 day and 10^th^ day of treatment relative to the group fed on the chow diet, results are mean ± SE, *** p<0.001 one-way ANOVA, Tukey’s multiple comparison test, n=6 each group comparing blood glucose between chow diet and high fat diet -fed group; **(ii)** Comparative fasting blood glucose in DIO mice upon exendin-4 administration at 0 day and 10^th^ day of treatment;, *** p<0.001, one-way ANOVA, Tukey’s multiple comparison test n=6, comparing fasting blood glucose (mean ± SE) at 0 day versus the 10^th^ day**;(iii)** Comparative fasting blood glucose in DIO mice upon I-M-150847 administration at 0 day and 10^th^ day of treatment; *** p<0.001, one-way ANOVA, Tukey’s multiple comparison test n=6, comparing fasting blood glucose (mean ± SE) at 0 day versus the 10^th^ day; **(iv)** Fasting blood sugar in I-M-150847 treated mice as compared to exendin-4 treated group and the group receiving normal saline as vehicle, results are expressed as mean ± SE; *** p<0.001, one-way ANOVA, Tukey’s multiple comparison test, n=6 for each group, comparing fasting blood glucose on the 10^th^ day between control mice receiving normal saline versus exendin-4 treated and I-M-150847 treated group. **C.** Intraperitoneal glucose tolerance (IPGTT) in DIO mice receiving normal saline as the vehicle (control), exendin-4 or I-M-150847 100nmol/kg bodyweight twice daily through subcutaneous administration, the data are expressed as mean ± SE ***p<0.001, *p< 0.05,(exendin-4 versus I-M-150847 treated group) as assessed by two-way ANOVA, Bonferroni (n=6, for each group) at different time points; glucose lowering during IPGTT as assessed by the area under the curve (AUC, mean ± SE, arbitrary units) in the vehicle, exendin-4 and I-M-150847 treated group, ***p<0.001, one-way ANOVA, Tukey’s multiple comparison test, n=6 each group comparing the vehicle, exendin-4, and I-M-150847 treated mice. **D**. Insulin tolerance test (ITT) in DIO mice receiving normal saline as the vehicle (control), exendin-4 or I-M-150847 100 nmol/kg body weight twice daily through subcutaneous administration, data are expressed as mean ± SE, **p<0.01, (exendin-4 and I-M-150847 treated group) assessed by two-way ANOVA, Bonferroni posttests (n=6, for each group) at different time points; ns= nonsignificant; glucose lowering upon insulin administration assessed by the area under the curve (AUC, mean ± SE, arbitrary units) in the vehicle, exendin-4 and I-M-150847 treated group, ***p<0.001, one-way ANOVA, Tukey’s multiple comparison test, n=6 each group comparing AUC of ITT in the vehicle, exendin-4 and I-M-150847 treated mice. **E.** Cumulative food intake in mice treated with vehicle, exendin-4, and I-M-150847 measured every day; the results are expressed as mean ± SE, *p<0.05 one-way ANOVA, Tukey’s multiple comparison test, n=6 for each group; *p<0.05, ***p<0.001, the statistical significance of the difference in cumulative food intake (I-M-150847 versus exendin-4 treated group, mean ± SE, n=6 for each group) at specific time points as assessed by two-way ANOVA, Bonferroni posttests. **F.** Body weight gain in mice receiving subcutaneous administration of normal saline (vehicle), exendin-4 or I-M- 150847 (n=6 each group). Results are expressed as mean ±SE, ***p<0.001 one-way ANOVA, Tukey’s multiple comparison test comparing % body weight gain among different groups. **p<0.01, ***p<0.001, the % bodyweight gain at specific time points (I- M-150847 versus exendin-4) was analyzed by two-way ANOVA, Bonferroni posttests (n=6 per group). **G.** Tissue weight of mice treated with normal saline (vehicle), exendin- 4, and I-M-150847 (n=6 per group), (i) liver, (ii) inguinal WAT, (iii) epididymal WAT, (iv) retroperitoneal WAT and (v) mesenteric WAT. Results are expressed as mean ±SE; **p<0.01, ***p<0.001, one-way ANOVA, Tukey’s multi comparison test (n=6 for each group).

Based on the evidence of enhanced glycemic benefits, we explored whether I-M-150847 provides appreciable weight loss in DIO mice. As **Fig 4E** shows, I-M-150847 causes a significant decrease in cumulative food intake compared to vehicle control (*p<0.05, one-way ANOVA, Tukey’s multiple comparison test, n=6 for each group) after 16 days of treatment. The decrease in cumulative food intake upon I-M-150847 treatment is significant compared to exendin-4 from the 7^th^ day of treatment (*p<0.05, two-way ANOVA, Bonferroni post-tests) that maximizes on the 16^th^ day of the study paradigm **(Fig 4E, \*\***\*p<0.001, two-way ANOVA, Bonferroni post-tests). Associated with the decreased food intake, we observed appreciable weight loss in I-M-150847 treated group that is significantly superior compared to the group treated with exendin-4 or vehicle **(Fig 4F,** ***p<0.001, one-way ANOVA, Tukey’s multiple comparison test, n=6 for each group). Further, I-M-150847 treatment causes a significant reduction in liver weight compared to exendin-4 mediated therapy and vehicle control (1.712 ± 0.05 grams in I-M-150847 treated group versus 2.47± 0.08 grams in the control group receiving normal saline as the vehicle, *** p<0.001, one-way ANOVA, Tukey’s multiple comparison test, n=6 for each group and 2.027± 62.37 grams in exendin-4 treated group p<0.05, one-way ANOVA Tukey’s multiple comparison test, n=6 for each group. **Fig 4G(i)**). We observed a significant reduction of inguinal WAT both in I-M-150847 and exendin-4 treated mice (152.7±27.94 grams and 140 ±14.96 grams in respective I-M- 150847 treated and exendin-4 treated groups compared to 337 ±0.45 gram in DIO mice receiving normal saline as vehicle control, ***p<0.001, one-way ANOVA, Tukey’s multiple comparison test, n=6 for each group). Similarly, there is a significant reduction in epididymal WAT (341.8±100.5 grams and 431.8 ±42.80 grams in I-M-150847 treated and exendin-4 treated group compared to 930.3 ±44.40 grams in the control DIO mice receiving normal saline as the vehicle,***p<0.001, one-way ANOVA, Tukey’s multiple comparison test, n=6 for each group), retroperitoneal WAT(68.0± 16.71 grams and 127.8 ± 15.72 grams upon respective I-M-150847 and exendin-4 treatment compared to 259.2±27.93 grams in control DIO mice receiving normal saline as vehicle ***p<0.001 and **p<0.01, one-way ANOVA, Tukey’s multiple comparison test, n=6 for each group) and mesenteric WAT (40.5± 3.905 grams and 54.67± 6.307 grams upon respective I-M- 150847 and exendin-4 treatment compared to 98.33± 9.2 grams ***p<0.001 and **p<0.01, one-way ANOVA, Tukey’s multiple comparison test, n=6 for each group), thereby demonstrating prominent reduction of adiposity both in I-M-150847- treated and exendin-4 treated DIO mice. The data, taken together, highlight the efficacy of GLP-1R GIPR dual agonist I-M-150847 in achieving improved glucose homeostasis and enhanced insulin sensitivity associated with the significant reduction of body weight, food intake, and adiposity in the DIO rodent model.

## Discussion

Multi-receptor agonism of incretins has recently emerged as a preferred therapy to achieve superior glycemic control and reduction of obesity (Gimeno et al., 2020; Samms et al., 2021). The concept of unimolecular GLP-1R and GIPR dual agonism first described by Finan et.al demonstrated the efficacy of hybrid peptides in reducing diabesity more effectively than the conventional GLP-1R agonists (Finan et al., 2013). Consistent with these findings, the dual GLP-1R, and GIPR agonist Tirzepatide (LY3298176) developed by Eli Lilly revealed better therapeutic ability than the most potent GLP-1R agonist semaglutide in reducing HbA1c, body weight, and serum triglycerides in SURPASS phase 3 clinical trials (Frias et al., 2021; Khoo and Tan, 2021; Min and Bain, 2021; Rosenstock et al., 2021; Slomski, 2021) thereby promising superior long-term glycemic control and significant reduction of obesity (Coskun et al., 2018), (Baggio and Drucker, 2021; Bailey, 2021).

The majority of the unimolecular incretin dual agonists reported to date were designed by intermixing GLP-1 and GIP amino acids in a single peptide using either glucagon (Finan et al., 2013), oxyntomodulin ((Bhat et al., 2013; Muppidi et al., 2016),(Yang et al., 2020), (Zhao et al., 2019), or GIP as the base peptide (Willard et al., 2020). The incretin receptor dual agonist I-M-150847 was designed based on the structure-based activation model of the Class B GPCRs (Hoare, 2005) taking clues from the conformational rearrangement upon ligand binding that initiates the second messenger signaling (Hoare, 2005; Parthier et al., 2007; Runge et al., 2008). The data on the crystal structure of GLP-1, exendin-4, and GIP bound to respective GLP-1R and GIPR extracellular domain (ECD) (Parthier et al., 2007; Runge et al., 2008; Underwood et al., 2010) provided us the critical insights for the designing of the middle and the C terminal region of the hybrid agonist. We focused on the Lys^30^ of GIP that aligns with the corresponding amino acid residue of oxyntomodulin and experimentally determined the activity by cAMP reporter assays. We replaced the tryptophan cage (Trp-cage) at the C terminus of exendin-4 (Lee et al., 2018; Neidigh et al., 2001; Runge et al., 2008; Wolff et al., 2021) despite its presence in Tirzepatide (Kegg Drug D11360) and stapled GLP-1R dual agonist O14(Yang et al., 2020) as previous studies reported the redundancy of the motif for the preservation of high binding affinity for GLP-1R (Lee et al., 2018).

We initiated structure-activity relationship (SAR) analysis with the synthesis of I-M- 150845 (**Table 1, Figure 1 -Figure supplement 1**) upon the addition of GLP-1 and GIP-specific amino acids to the first 9 amino acid residues of the peptide I-M-150844 that corresponds to the N-terminal sequence of exendin-4. I-M-150845 showed dual GLP-1R and GIPR activity in cAMP reporter assays (**Table 1**). From the structural perspective, the amino acid residues Phe^22^, Val^23^, Leu^26^, and Leu^27^ of the amphipathic alpha-helix of GIP that forms-minute hydrophobic interactions with GIPR ECD are present in the hybrid peptide I-M-150845. In addition, we retained Asp^15^, Gln^19^, Gln^20^, Leu^26^, and leu^27^, the amino acids that form an extensive hydrogen-bonding network with several residues of GIPR ECD in I-M-150845 to provide GIPR-specific activity. In contrast, the amino acids of the exendin-4/GLP-1 amphipathic alpha-helix that support the hydrophilic interactions with GLP-1R ECD are absent in I-M-150845. Of the 7 amino acids of exendin-4, Ala^18^, Val^19^, Phe ^22^, Ile ^23^, Trp^25^, Leu^26^, and Pro^31^ that form hydrophobic interaction with GLP-1R ECD, only Phe^22^ and Leu^26^ are retained in hybrid dual agonist I-M-150845. We hypothesized that Val^27^ at the I-M-150845 has a compensatory influence as it establishes hydrophobic contact as well as hydrogen bonding with GLP-1R ECD in the receptor-ligand crystal complex (Underwood et al., 2010). To experimentally ascertain the importance of Val^27^ for GLP-1R specific activity, we synthesized a new incretin agonist I-M-150846 where the C-terminal amino acids Val^27^-Gly^31^ of I-M-150845 are substituted with 11-amino acid C terminus (Met^27^-Ala^37^) of oxyntomodulin (**Figure 1 Figure supplement 2**). The substitution removes Val^27^ but introduces Lys^30^ which in GIP establishes an intricate hydrogen-bonding network with several residues in the C terminal part of GIPR ECD (Parthier et al., 2007). The incretin agonist I-M-150846 revealed a complete loss of function at GLP-1R while retaining GIPR activity **(Table 1).** We then replaced GIP-specific Tyr^10^-Asn^24^ of I-M-150846 with amino acid residues Leu^10^- Glu^24^ of exendin-4 (Figure **1- Figure supplement 3**) that forms superior helical architecture with GLP-1R ECD (Runge et al., 2008). The resultant incretin agonist I-M-150847 showed robust GLP-1R and GIPR dual agonism signifying that in the absence of Trp cage, the presence of the amino acid residues Leu^10^- Glu^24^ of exendin-4 in the hybrid agonist peptide is critical to establish a stable interaction with the GLP-1R ECD thereby restoring the functional activity towards GLP-1R as measured by cAMP reporter assay. The data also implies that Phe^22^, Leu^26^, and Lys^30^ are the crucial amino acids that confer binding to the GIPR ECD required for the GIPR-specific activity of I-M-150847.

Variable propensities of internalization and recycling of GLP-1R determine the potential of GLP1-RAs in delivering insulinotropic response (Jones et al., 2018). We first reported sustained cAMP signaling following prolonged internalization of GLP-1R upon treatment of BRIN-BD11 pancreatic beta cells with GLP-1Tmr, exendin-4, and liraglutide (Kuna et al., 2013),(Girada et al., 2017),(Bele et al., 2020). Endocytic sorting of GLP-1R to lysosome upon treatment with GLP-1Tmr and exendin-4 was reported in cultured pancreatic beta cells(Kuna et al., 2013),(Jones et al., 2018). In contrast, exendin F1, an exendin -4 analog, elicits a fast-recycling response of GLP-1R and provides better glycemic benefits compared to other GLP-1R monoagonists (Jones et al., 2018). In our present study, we observed prompt redistribution of functional GLP-1R at the plasma membrane following I-M-150847 treatment in contrast to exendin-4 -mediated prolonged receptor internalization. Our data shows that the I-M-150847- mediated cAMP generation is discrete as we observe an abrupt attenuation of receptor-mediated cAMP accumulation 30 minutes after the treatment with I-M-150847 which is in contrast to the exendin-4 mediated activation where continuous cAMP accumulation is observed till an hour after the agonist exposure. In this context, the gain of cAMP generating response following agonist rechallenge that is specific for I-M-150847- mediated activation is indicative of reduced internalization and or faster GLP-1R recycling that may account for the higher distribution of the sensitized receptor at the plasma membrane following I-M- 150847 treatment.

Functional consequences of the preservation of incretin receptors in a resensitized state are presently unclear. In our study, we observe superior insulinotropic activity upon acute treatment with I-M150847 compared to exendin-4. However, GSIS is blunted in the case of chronic exposure to exendin-4 or I-M-150847 signifying homologous desensitization of the incretin receptor both in the case of prolonged internalization in exendin-4 - treated cells as well as upon faster recycling or reduced internalization that we observe upon I-M-150847 treatment. In contrast, in the case of mouse adipocytes that express GIPR (McIntosh et al., 2012), we observe retention of insulin sensitivity as well as cAMP accumulation response after chronic exposure to I-M-150847.GIPR internalization and desensitization upon treatment with the cognate agonist have been earlier documented (Ismail et al., 2015). In this context, the functional response of I-M- 150847 in mouse adipocytes reveals retention of GIPR in a sensitized state upon chronic exposure to the agonist supporting the downstream signaling cascade that regulates the cellular metabolism. The data is also indicative of the cell type variability of incretin receptor desensitization as has been reported in the case of the M3 muscarinic receptor(Torrecilla et al., 2007), (Kelly et al., 2008).

The efficiency of I-M-150847 in eliciting insulin sensitivity in adipocytes translates into better in vivo glycemic benefits as we observe significantly improved reduction of fasting blood glucose and better glucose tolerance in DIO mice upon I-M-150847 treatment. We also observed a significant reduction of food intake and weight loss in I-M-150847 treated mice relative to the group administered with exendin-4 or vehicle. However, whether this greater anorexic response upon I-M-150847 treatment is a consequence of the preservation of GLP-1R and or GIPR at a sensitized state at the feeding centers in the hypothalamus deserves attention in future research.

A key question regarding incretin receptor dual agonism is the contribution of GIPR in the attainment of superior glycemic control. Several reports highlight the benefits of LAGIPRA in enhancing insulin sensitivity through activation of the Akt signaling pathway that is associated with the storage and oxidation of glucose in adipose tissues, (Samms et al., 2021; Samms et al., 2020). However, similar attributes of improved glucose tolerance and insulin sensitivity have been reported in contrasting mouse models representing GIPR overexpression (Kim et al., 2012) and adipose tissue-specific GIPR knockout (Joo et al., 2017). Our data with 3T3-L1 adipocytes demonstrate a significant increase in insulin-stimulated glucose uptake upon chronic I-M-150847 treatment despite the increase in the cAMP generation which is indicative of increased lipolysis. The data is in agreement with the previous report of increased lipoprotein lipase activity upon GIP treatment in cultured preadipocytes (Eckel et al., 1979) as well as the enhancement of insulin sensitivity in epididymal WAT upon LAGIPRA treatment in DIO mice due to increased Akt phosphorylation (Varol et al., 2014). Thus, while it is likely that GIPR knock-out mice showed enhanced insulin sensitivity due to reduced GIPR - mediated lipolysis and consequent less free fatty acid (FFA) generation, GIPR overexpression or treatment with LAGIPRA or GLP-1R GIPR dual agonists may augment insulin sensitivity through activation of metabolic pathways that are associated with glucose and FFA metabolism in the adipose tissue. We note that a detailed mechanistic study is required to fully test the hypothesis.

Collectively, the preclinical data presented in this study demonstrate I-M-150847 as a novel GLP-1R and GIPR dual agonist that promotes expeditious resensitization of GLP- 1R and preserves the functional attributes of GIPR upon chronic exposure in adipocytes thereby augmenting better insulinotropic response and enhancing insulin sensitivity consequentially improving the systemic glycemic control. Further, I-M-150847 reduces food intake, ensures significant body weight loss, and decreases adiposity highlighting the therapeutic potential of the novel hybrid agonist in ameliorating T2DM and obesity.

## Materials and Methods

### Key resources table

Following reagents are used in this study.

| REAGENT | SOURCE | IDENTIFIER |
| --- | --- | --- |
| CHEMICALS, PEPTIDES, AND RECOMBINANT PROTEINS |  |  |
| RPMI 1640 HiGlutaXL | Himedia | AL-060G |
| DMEM HIGH GLUCOSE | Himedia | AL-219A |
| FBS Heat Inactivated (US ORIGIN) | Himedia | RM-10409 |
| TRYPsin EDTA (10X) | Himedia | TCL-034 |
| Penicillin Streptomycin solution (100X) | Himedia | A001 |
| Gentamycin | Himedia | A005 |
| DMSO | SIGMA | M81802 |
| LIPOFECTAMINE 3000 | THERMO | L3000001 |
| IBMX | SANTA CRUZ | SC-201188 |
| GLUCOSE | SIGMA | G-8270 |
| PHOSPHATE BUFFER SALINE | INVITROGEN | 70011044 |
| BETA MERCAPTOETHANOL | SIGMA | M6250 |
| RNAiso Plus | Takara | 9108 |
| BOVINE SERUM ALBUMIN | SIGMA | A2153 |
| FAT-FREE BSA (FRACTION V) | SIGMA | 5479 |
| STEADYLITE Plus | PERKIN ELMER | 6066751 |
| ATP | SIGMA | A7699 |
| PROTEASE INHIBITOR COCKTAIL | CALBIOCHEM | 539131 |
| LIRAGLUTIDE (VICTOZA) | NOVO NORDISK | Acylated GLP-1R agonist |
| PHOSPHATASE INHIBITOR | CALBIOCHEM | 524625 |
| RNA Later solution | AMBION | AM7020 |
| Triton-X 100 | SIGMA | T8787 |
| Dr. Trust Glucometer and strips | Dr. Trust USA | 9001 |
| IN-M-150845 | Designed in-house,<br>synthesized, and, purified by<br>GL BIOCHEM, CHINA | Procured through Biotech<br>Desk India |
| IN-M-150846 |  |  |
| IN-M-150847 |  |  |
| Exendin F1 | Dr. Ben Jones, Imperial College, London |  |
| Oxyntomodulin | Tocris | 2094 |
| GIP | Genscript | I |
| 2-nitro-phenyl beta galactopyranoside | SIGMA | N1127 |
| Paraformaldehyde | SIGMA | 158127 |
| Vectashield | Vector Labs | H-1200-10 |
| Plasmid GLP-1R GFP | Mitra Lab | Kuna et.al |
| Plasmid CRE Luciferase | Richard Day, Indiana University | Kuna et.al 2013(Kuna et al., 2013) |
| Plasmid GIPR | Bernard Thorens | ADDGENE#14942(Gremlich et al., 1995) |
| Plasmid GCGR | Brian Finan | Novo Nordisk |

| CRITICAL COMMERCIAL ASSAYS |  |  |
| --- | --- | --- |
| BCA PROTEIN ASSAY KIT | SANTA CRUZ | sc-200629 |
| INSULIN ELISA KIT (RAT/MOUSE) | MERCK | EZRMI-13K |
| cAMP ELISA kit | ENZO | ADI-900-067 |
| TB Green® Premix Ex Taq™ II | Takara | RR820B |
| PRIMESCRIPT cDNA synthesis kit | Takara | RR014B |
| PLASMID ISOLATION KIT | INVITROGEN | K210005 |

| OLIGONUCLEOTIDES |  |
| --- | --- |
| RT-qPCR Primers | IDT |

| EXPERIMENTAL MODELS/ CELL LINES |  |  |
| --- | --- | --- |
| BRIN-BD11 CELL LINE | ECACC | 10033003 |
| HEK-293T CELL LINE | ATCC | CRL-3216 |
| 3T3L1 preadipocytes | ATCC | CL-173 |
| C57BL/6 MICE | Jeeva Life Sciences, Hyderabad | JLS-000664 |
| Animal Feed (HIGH FAT DIET/CHOW DIET) | NIN, Hyderabad |  |

## Methods

### Peptide Synthesis

Synthesis of the 11- amino acid residue peptide I-M-150844 was carried out in Biotage Syro I automated peptide synthesizer using Wang resin and following Fmoc chemistry. The amino acid that was first loaded to the resin is Phenylalanine and following installation, the capping of residual functional groups on the resin was carried out by treatment with 33% acetic anhydride and pyridine. Before loading of next amino acid, the Fmoc group of the amino acid was deprotected through treatment by 40% piperidine / DMF and the incorporation of the next amino acid was carried out using coupling reagent HBTU (0.5 mmol) and DIPEA/NMP (2 mmol). The peptide was cleaved from the resin by mixing it with TFA/H_2_O/TIPS (95:2.5:2.5, 4 ml) and TFA was removed by evaporation on a rotatory evaporator. The crude mixture was triturated (diethyl ether) and purified by preparative HPLC using Agilent column ZORBAX 300SB-C18 (5 μm, 9.4*250 mm).

The other peptides I-M-150845, I-M-150846, and I-M-150847 were designed in-house and outsourced for chemical synthesis and purification to GL Biochem China. The C terminal amidated version of I-M-150847 was synthesized by Aurigene Pharmaceutical Services, India.

### Circular dichroism spectroscopy

Determination of the secondary structure of I-M-150847 was carried out by using Circular Dichroism Instrument (JASCO, J1500, Easton MD) (Kelly et al., 2005; Waterhous and Johnson, 1994). The spectra from 200 to 260 nm were taken with a data interval of 0.5nm. A bandwidth of 1.00nm with a 100 nm/min scanning rate was used to determine the property.10µM of I-M-150847 was used for the study and 50mM Phosphate buffer serves as the blank background. The data were represented as Ellipticity (mdeg) vs wavelength (nm) at the Y and X-axis respectively.

### Cell Culture

Pancreatic beta cells, BRIN-BD11 (cat no.10033003) were procured from the European Collection of Authenticated Cell Cultures. The cells were cultured in RPMI media 1640 HiGlutaXL with 10% heat-inactivated fetal bovine serum (US origin) supplemented with 1mM sodium pyruvate, 50µM β-mercaptoethanol,10mg/mL gentamycin, 100U/mL penicillin, and 100mg/mL streptomycin at an incubation temperature of 37⁰ C with 5% CO_2_ following the method of Kuna et. al (Kuna et al., 2013). Human Embryonic Kidney cell lineage (HEK 293T) cells were procured from ATCC (CRL3216). The cells were cultured in DMEM-Hi Glucose media supplemented with 10% heat-inactivated FBS, 10mg/mL gentamycin, 100U/mL penicillin, and 100mg/mL streptomycin at an incubation temperature of 37⁰ C with 5% CO_2_, cultured following the protocol recommended by ATCC. 3T3 L1 mouse preadipocytes (ATCC (CL-173)) were sub-cultured using DMEM supplemented with 10% newborn calf serum and 100U/mL penicillin, and 100mg/mL streptomycin at an incubation temperature of 37⁰ C with 5% CO_2_ following ATCC protocol. The preadipocytes were differentiated into adipocytes following the protocol of Mitra et.al (Mitra et al., 2004)

### GLP-1R, GIPR, GCGR activation

Each peptide was tested for its ability to activate GLP-1R, GIPR, or GCGR in a cell-based cAMP-responsive element luciferase reporter assay following the method of Kuna et.al (Kuna et al., 2013). Briefly, HEK 293T cells were cultured in a 60mm tissue culture treated dish. Upon 70% confluency, the cells were transiently transfected with cAMP-responsive element luciferase reporter plasmid, beta-galactosidase plasmid, and GIP-R or GLP-1 R, or GCGR plasmid in a 3:1:3 ratio. The transient transfection was carried out using lipofectamine 3000 following the manufacturer’s protocol. After 4 hours of transfection, the cells were seeded in a tissue culture grade 96 well plate at a density of 50000 cells / well. After overnight incubation, the cells were treated with different concentrations of agonists for the different cell-based assays. After 4 hours of treatment, the media was aspirated and cells were lysed and the luminescence was measured using Steady lite Plus reagent (Perkin Elmer Life and Analytical Science, Waltham, MA) after 15 minutes of shaking at 300 rpm at room temperature in dark. 2- nitro-phenyl beta galactopyranoside (Sigma) was used to carry out a beta-galactosidase assay for correcting the inter-well transfection variability.

### cAMP accumulation assay

The accumulation of cAMP was measured in cultured BRIN-BD11 at different time points upon agonist treatment. 1µM I-M-150847 and exendin-4 were tested for the time-course of cAMP generation using a direct cAMP ELISA kit (Enzo Life Sciences, Cat. No. ADI-900-163). The assay was carried out following the protocol of Girada et. al (Girada et al., 2017) with modifications. Briefly, BRIN-BD11 cells were seeded in 24 well-treated plates at a density of 0.1 million cells/well. After overnight incubation, the cells were incubated again for 60 minutes in the KRB-BSA medium (with 1.1mM glucose) after which the cells were washed and treated with 1µM I-M-150847 or Exendin-4 for 3 minutes, and then the treatment medium was washed off and replaced with KRB-BSA medium with 200µM IBMX and 5.5mM glucose and incubated for 1, 10,30, and 60 minutes after which the cells were lysed and processed for the cAMP ELISA assay. Protein quantification was carried out using BCA following kit protocol.

To assess chronic cAMP generation in 3T3-L1 adipocytes, the cells were treated with 1µM I-M-150847 or exendin-4 overnight in serum-free media following which they are washed and incubated with KRB buffer with 0.1%BSA containing 1.1mM glucose. The cells were then exposed to a treatment medium comprising fresh KRB-BSA buffer with 5.5mM glucose, 200µM IBMX, and 1µM I-M-150847 or exendin-4 for 3 minutes, and following wash incubated in KRB buffer with 5.5mM glucose and 200µM IBMX for 30 minutes. The cells were then lysed and processed for cAMP ELISA assay. Protein quantification was carried out using BCA following the kit protocol.

### Agonist rechallenge assay

Agonist rechallenge was carried out following the protocol by Jones et.al (Jones et al., 2018) with modifications. Briefly, BRIN-BD11 cells were seeded in 24 well tissue culture treated plates at a density of 0.1 million cells/ well. After overnight incubation in the complete medium, the cells were incubated for 60 minutes in the KRB-BSA medium with 1.1mM glucose after which the cells were washed and treated with 1µM I-M- 150847 or Exendin-4 in a KRB-BSA medium containing 200µM IBMX and 5.5mM glucose for 3 minutes. The treatment medium was then washed off and replaced with a fresh KRB-BSA medium containing 5.5mM glucose and 200µM IBMX and incubated for 45 minutes. The cells were re-challenged with agonists for 10-minutes followed by lysis and subsequently processed for the cAMP ELISA assay. Protein quantification was carried out using BCA following kit protocol.

### GLP-1R internalization and post-endocytic sorting

Assessment of GLP-1R internalization was carried out in BRIN-BD11 pancreatic beta cells and monitored using confocal microscopy. The cells were transfected with GLP- 1R-GFP using lipofectamine 3000 and seeded in six-well plates on 22*22mm sterile cover glass coated with poly-L-lysine. 48 hours after transfection, the cells were washed with sterile PBS and incubated with I-M-150847 (1µM) or Exendin-4 (1µM) in 200µL of Krebs-HEPES buffer for 60 min at 4°C in a humidified chamber in the dark. Cells were then washed in PBS and incubated at 37°C for 1 min, 10 minutes, 30 minutes, and 60 minutes after which they are fixed in 4% paraformaldehyde and imaged using a Zeiss LSM 880 confocal laser scanning microscope (Carl Zeiss, Oberkochen, Germany) equipped with HXP 120V light sources. Pinhole diameter was maintained at 1 airy unit. Fields were selected using a 20X objective and image acquisition was carried out using a 63 X oil immersion objective lens.

The post-endocytic sorting of GLP-1R after activation was carried out following the method of Kuna et.al with modifications (Kuna et al., 2013). Briefly, BRIN-BD11 cells were transfected with GLP-1R-GFP, and 60 hours after transfection were treated with 1µM I-M-150847 or Exendin-4 and incubated at 4°C, in a dark humidified chamber for one hour. The cells were then washed with ice-cold PBS and treated with Lysotracker red (100nM) for 45 minutes in RPMI-1640 medium, washed with PBS, and fixed in 4% paraformaldehyde. The coverslips were mounted on glass slides by using Vectashield mounting media and the imaging was carried out in Zeiss LSM 880 confocal laser scanning microscope (Carl Zeiss, Oberkochen, Germany) equipped with HXP 120V light sources with a pinhole diameter of 1 airy unit. The emission was measured at 500- 550nm bandpass for GFP and 575-615 nm for lysotracker red. The images were acquired by using a 63X oil immersion objective lens with 2X optical zoom and analyzed using Zenlite 2011 program.

### Glucose- stimulated Insulin secretion

Insulin secretion was assessed using a Millipore rat/ mouse insulin ELISA kit (Merck, EZRMI-13K) following the method of Asalla et.al.(Asalla et al., 2016) Briefly, BRIN- BD11 cells were seeded in 24 well tissue culture plates at a density of 0.1 million cells / well. In the next day, the cells were treated with Kreb’s Ringer Bicarbonate buffer (KRB) [115 mM NaCl, 4.7 mM KCl, 1.28 mM CaCl_2_, 1.2 mM KH_2_PO_4_, 1.2 mM MgSO_4_, 10 mM NaHCO_3_, 0.1% (wt./vol) BSA, pH 7.4] for 1h without glucose. Glucose-stimulated insulin secretion (GSIS) was assessed in the presence of indicated concentrations of glucose and incretin receptor agonists for 30 min and the supernatant was collected, centrifuged, and processed for the ELISA as per the manufacturer’s protocol. The cells were lysed with 0.1% Triton-X 100 and processed for protein quantification to normalize the insulin secretion. The data were expressed as fold-over basal insulin secretion (pg insulin/µg protein). The insulin secretion at 0.1mM glucose is considered basal secretion.

For assessment of GSIS during prolonged agonist treatment, BRIN-BD 11 cells were treated with 1µM agonists in a 2% FBS-RPMI medium for 6 hours. The cells were then washed with KRB buffer and incubated in KRB-0.1%BSA for 1 hour. The incubation medium was replaced with a fresh KRB-BSA medium in the presence of 16.7mM glucose and 1µM agonists for 30 minutes following which the supernatant was collected for insulin ELISA and the cells were lysed to determine the protein to normalize the insulin secretion. 10 µL of the sample supernatant was used for the insulin ELISA assay. The quantity of insulin was measured as the picogram (pg) of insulin per milligram (mg) of protein and expressed as a fold increase over the basal insulin secretion. The insulin secretion at 0.1mM glucose is considered basal secretion.

### Glucose uptake assay

Glucose uptake was carried out on the 9^th^ day differentiated 3T3-L1 adipocytes with a fluorescent D- glucose analog 2-[N-(7-nitrobenz-2-oxa-1,3diazol-4-yl) amino]-2-deoxy-D glucose (NBDG) following the method of Zou et.al (Zou et al., 2005). Briefly, 3T3 L1 adipocytes were treated with 1µM of I-M-150847 or Exendin-4 overnight in serum-free DMEM. The cells were then washed and incubated with KRB buffer containing 0.1% BSA for 60 minutes. Post incubation, the cells were treated with 100nM of insulin for 30 minutes after which 150 µM of 2-NBDG was added for 20 minutes. The reaction was stopped using ice-cold PBS, and the cells were lysed with the lysis buffer containing 0.1% Triton X 100 in dark on an orbital shaker. The lysates were transferred to the Corning 96 well polystyrene Black microplate and read in triplicate at 485/535 nm excitation/emission in a multi-mode plate reader.

### Total RNA isolation, cDNA preparation, and qRT PCR

The RNA extraction was carried out by phenol-chloroform extraction protocol from cultured BRIN-BD11 pancreatic beta cells.1 µg of total RNA was used for first-strand cDNA synthesis. Takara Prime script cDNA synthesis kit was used to carry out the two-step cDNA synthesis following the manufacturer’s protocol. The mRNA for GLP-1R and GIPR was amplified and quantified by quantitative real-time PCR using Quant Studio 5 (A&B Biosystems). 2XSYBR master mix (Takara Life Sciences) reagent was used for the assessment of the relative abundance of mRNAs measured using the 2^-ΔΔCT^ method (Livak and Schmittgen, 2001). Normalization was accomplished using GAPDH as an invariant reference. Primers used for amplification are as follows: GLP1R *f*: TATTGGCTCATCATACGCTTG; GLP1R*r:* GTCTGCATTTGATGTCGGTCT; GIPR *f*: CGAAGTCAAAGCCATTTGGT; GIPR *r*: CCAGCCTTAGTCGGTAGTCG; Beta actin *f*: CCCATACCCACCATCACACC; Beta actin *r*: GAGAGGGAAATCGTGCGTGAC.

### Animal Experiments

Animal experiments were performed following the guidelines of Control and Supervision of Experiments on Animals (CPCSEA) upon permission from the Institutional Ethical Committee Hyderabad, India (UH/IAEC/KS/2021-1/36).C57BL/6 male mice of 5 weeks of age were procured from Jeeva Life Sciences, Hyderabad (Cat No: JLS-000664). Only male mice were used for the study since female mice are mostly protected from diet-induced obesity (Pettersson et al., 2012).

### Pharmacological and Metabolism studies

The mice were group-housed or single-housed on a 12:12 h light-dark cycle at 22°-24°C and were fed ad-libitum on a diabetogenic diet (Bele et al., 2020) which is a high-sucrose diet with 58% kcal from fat for 75 days. The mice were then randomized based on body weight, food intake, and blood glucose before the initiation of the study and were divided into 3 groups (n=6 / group): control receiving normal saline as a vehicle, comparator group receiving 100 nmol/kg body weight exendin-4 twice daily and test group receiving 100nmol/kg body weight I-M-150847 twice daily through subcutaneous administration for 4 weeks. Food intake and body weight were measured every day after the first injection of agonists. Fasted blood glucose was measured at the study initiation, on the 10^th^ day of treatment, and at the termination of the experiment.

#### Glucose tolerance test

On the 20^th^ day of treatment, DIO mice were subjected to fasting for 5 hours and following which the intraperitoneal injection of glucose (2g/kg body weight) (D-glucose [Sigma] 40% w/v in 0.9% normal saline) was administered. Glucose level was monitored by a glucometer upon collecting blood from the tail vein before (0) and after (15, 30, 60, 90, and 120 minutes) after glucose administration.

#### Insulin tolerance test

On the 24^th^ day of treatment, the animals fasted for 4 hours after which the animals received an intraperitoneal injection of insulin (0.75U/kg body weight, Humulin R, Eli Lilly, and Company). The blood from the tail vein was collected before (0) and after (15, 30, 60, and 90 minutes) insulin administration, and blood glucose was monitored using a glucometer.

#### Tissue weight measurement

For the measurement of tissue weight, animals were sacrificed on the 28^th^ day of treatment and mesenteric, retroperitoneal, inguinal, and epididymal WAT were collected. Blood and Liver were also collected during the sacrifice and kept at −80° C for further processing.

### Statistical analysis

Statistical analyses were performed with GraphPad Prism 6.0 and presented as means ± SEM. The analysis of the experiments was performed with one-way ANOVA (Tukey’s multiple comparison test) and Student’s t-test or two-way ANOVA Bonferroni posttests. *P* values lower than 0.05 were considered significant.

### Ethics

The animal studies were carried out following the guidelines of Control and Supervision of Experiments on Animals (CPCSEA) upon permission from the Institutional Ethical Committee Hyderabad, India (UH/IAEC/KS/2021-1/36).

## Author Contributions

RB and SB jointly carried out the experiments with the help of JE and equally contributed to the study. SK, TD, and AI provided the necessary support for conducting experiments. NC and VR synthesized some of the peptides and carried out physical characterizations. RB, SB, and PM carried out the in-silico analysis of the structure of novel incretin peptides. PM conceptualized the project and supervised the experiments, designed the peptides, analyzed and interpreted the data, wrote the manuscript upon discussion with RB and SB, and take the responsibility for the integrity and accuracy of the data. All authors have full access to the data related to the manuscript and approved the final version of the manuscript for publication.

## Acknowledgments

The work has been supported by Department of Science and Technology Grants; EMR/2016/007057 provided to PM, VR, and AI. Fellowship of RB and SB is supported by the Council of Scientific and Industrial Research (CSIR No: 09/0985 (0032)/2019-EMRI) and ICMR senior research fellowship (No 2017-3891-CMB/BMS). SB and RB are registered at the Manipal Academy of Higher Education (MAHE) for their doctoral degree. Exendin F1 is a generous gift from Dr. Ben Jones, Imperial College London.

---

**Figure1-Figure supplement 1:**
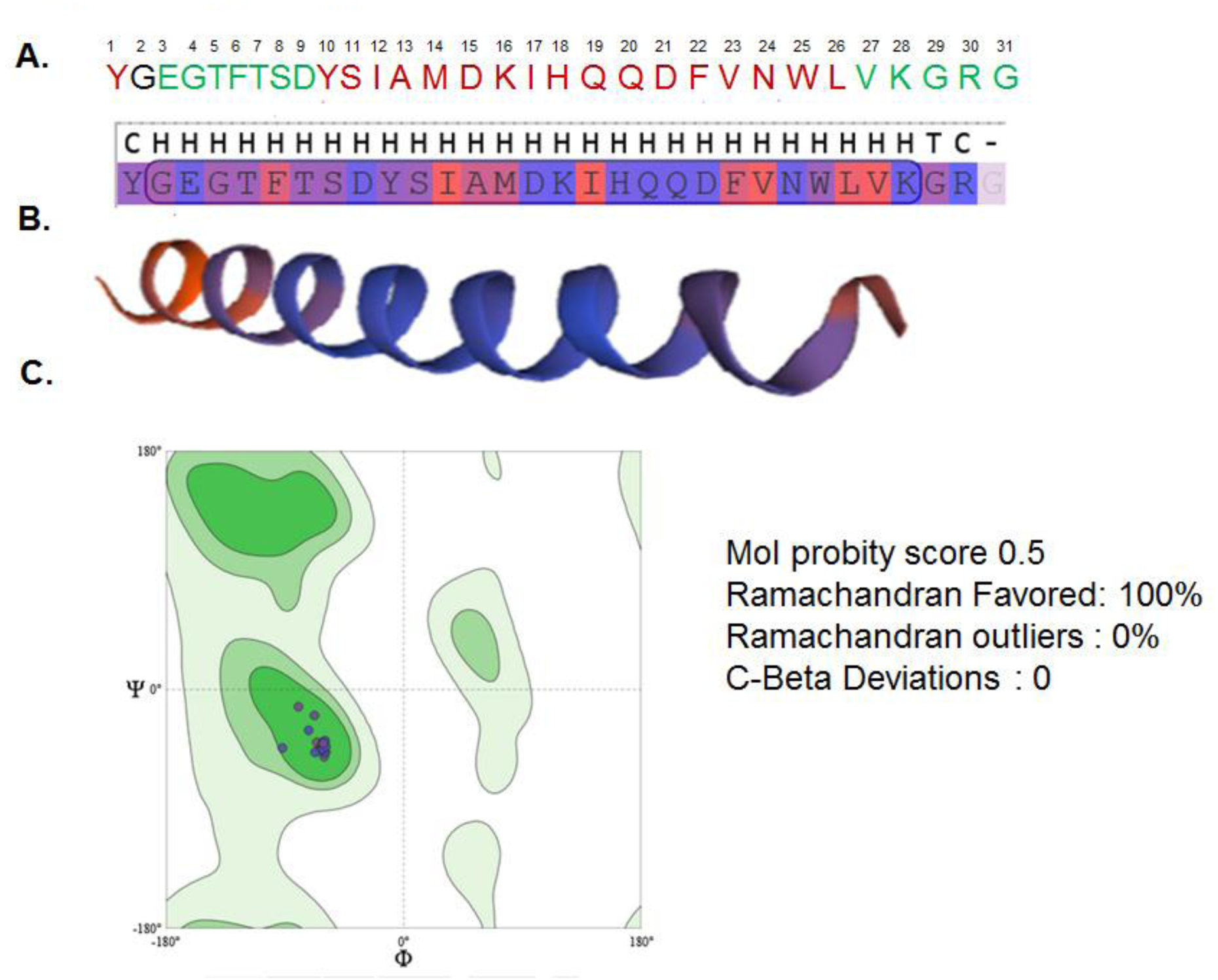
Design of the peptide I-M-150845. A. Amino acid constituents of peptide I-M-150845. The amino acids common to exendin-4, GLP-1, and GIP are colored black, green, and red respectively. B. Cartoon of the predicted secondary structure of I-M-150845 as modeled by Swiss-model protein structure homology modeling server. C=loop or irregular, H=α helix, T= Hydrogen bonded turns. C. Ramachandran plot for I-M-150845 using Mol Probity version 4.4 in Swiss-model protein structure homology modeling server.

**Figure1- Figure supplement 2:**
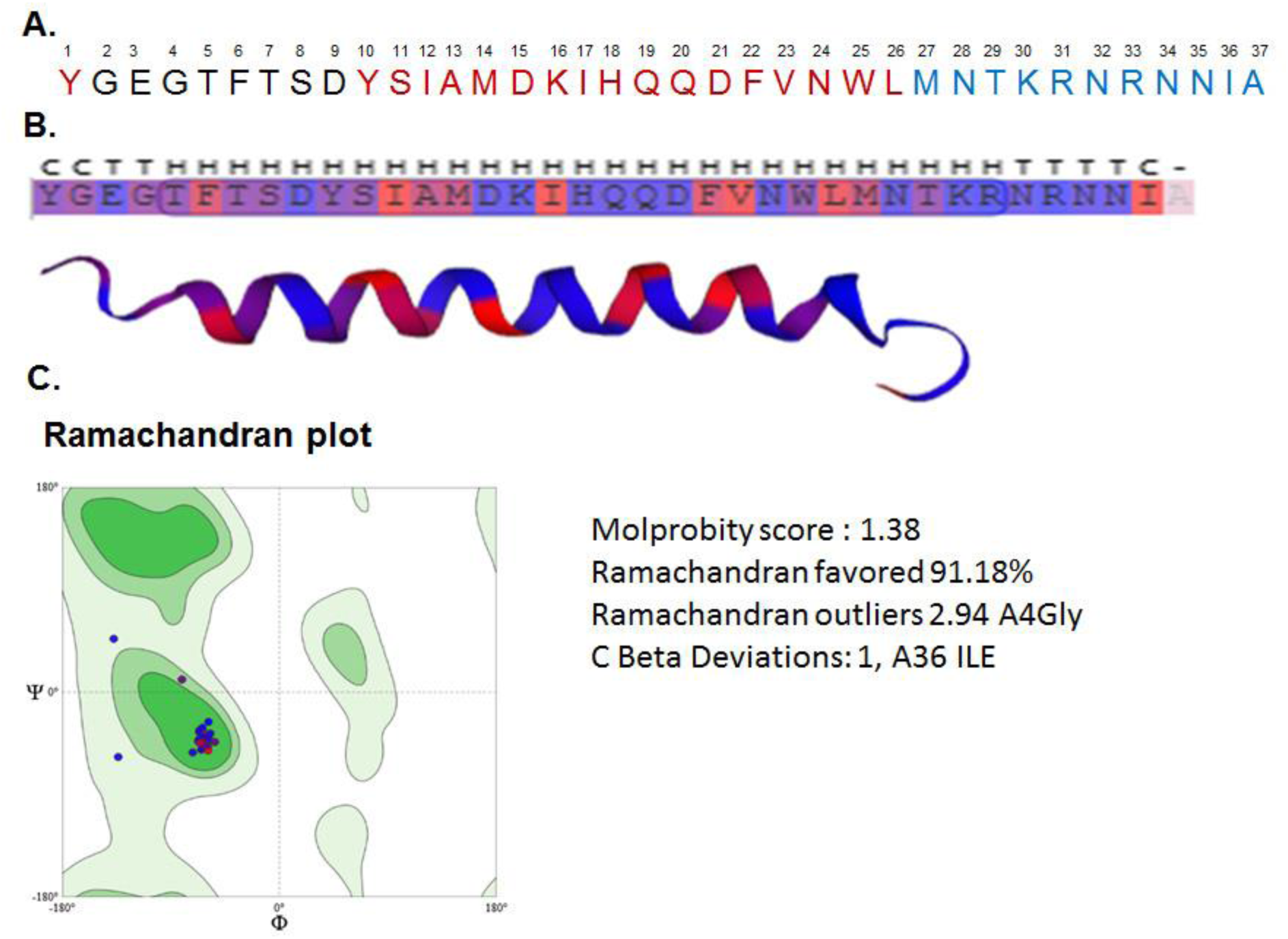
Design of the peptide I-M-150846. A. Amino acid constituents of peptide I-M-150846. The amino acids common to exendin-4, GIP, and oxyntomodulin are colored black, red, and blue respectively. B. Cartoon of the predicted secondary structure of I-M-150846 as modeled by Swiss- model protein structure homology modeling server. C=loop or irregular, H=α helix, T= Hydrogen bonded turns. C. Ramachandran plot for I-M-150846 using Mol Probity version 4.4

**Figure 1-Figure supplement 3:**
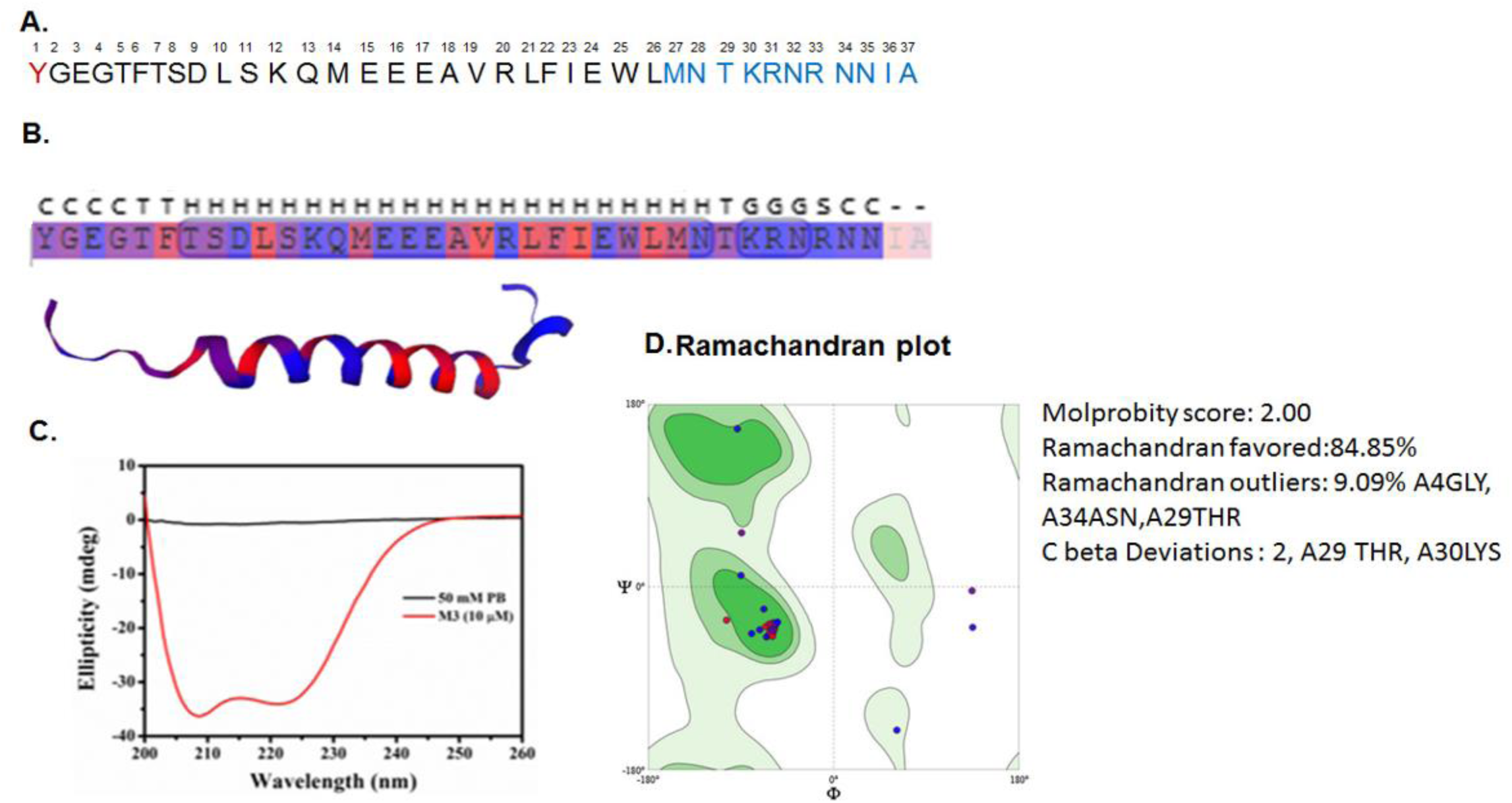
Design and physical characterization of peptide I-M-150847. A. Amino acid constituents of I-M-150847. The amino acids common to exendin-4, GIP, and oxyntomodulin are colored black, red, and blue respectively B. Cartoon of the predicted secondary structure of I-M-150847 as modeled by Swiss- model protein structure homology modeling server. C=loop or irregular, H=α helix, T= Hydrogen bonded turns. G= 3-helix (3_10_- helix), S= bend. C. CD analysis of I-M-150847. D. Ramachandran plot for I-M-150847 using Mol Probity version 4.4

**Figure-1 Figure supplement 4:**
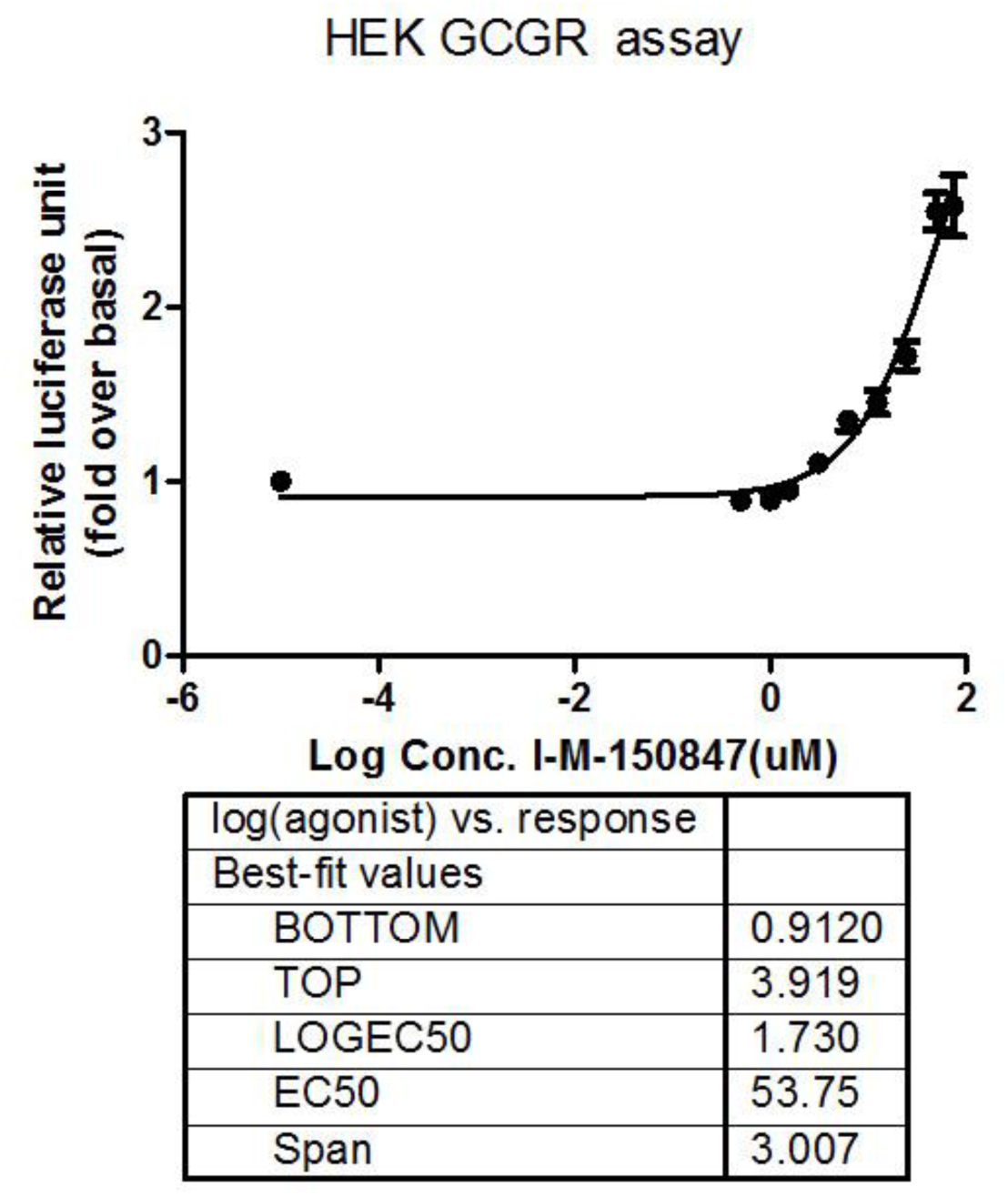
Dose response curve of I-M-150847 mediated cAMP response in HEK GCGR cell line measured by cAMP-responsive element luciferase reporter assay. Data are represented as mean ± SE of 3 independent experiments

**Figure 1 Figure supplement 5:**
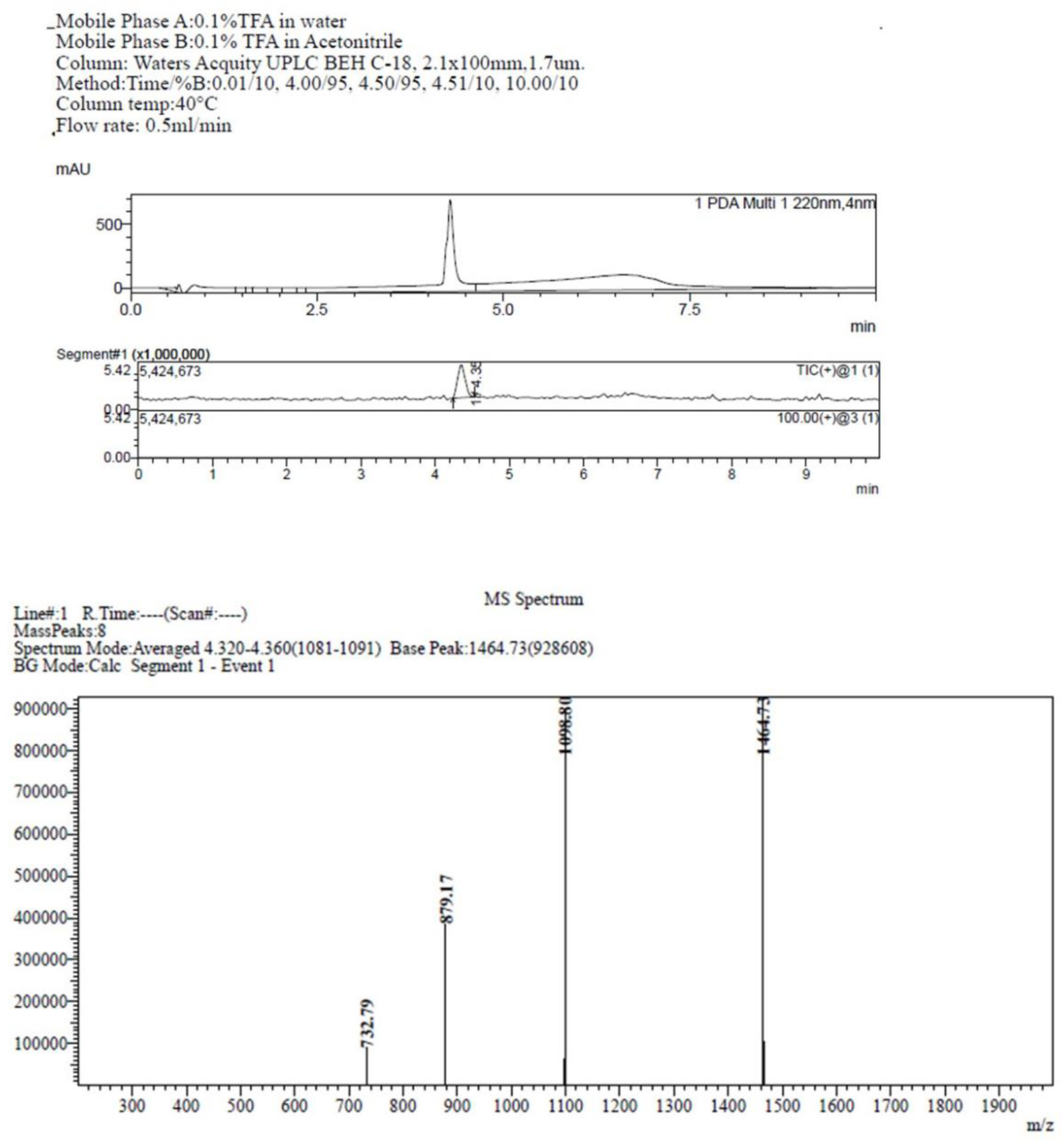
LC-MS analysis of peptide I-M-150847 amide.

**Figure 1-Figure supplement 6:**
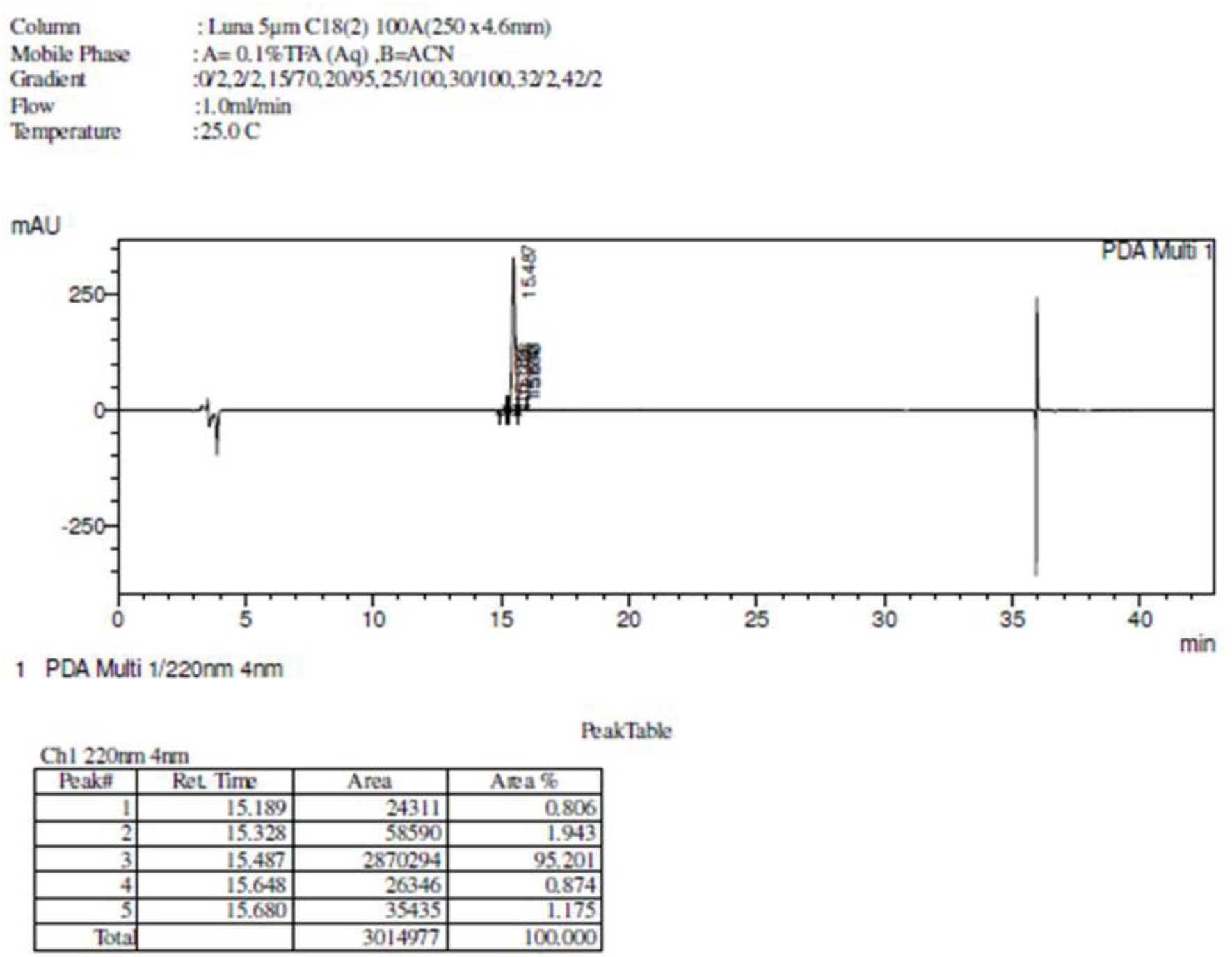
RP-HPLC of peptide I-M-150847 amide

**Figure -3 Figure supplement 1:**
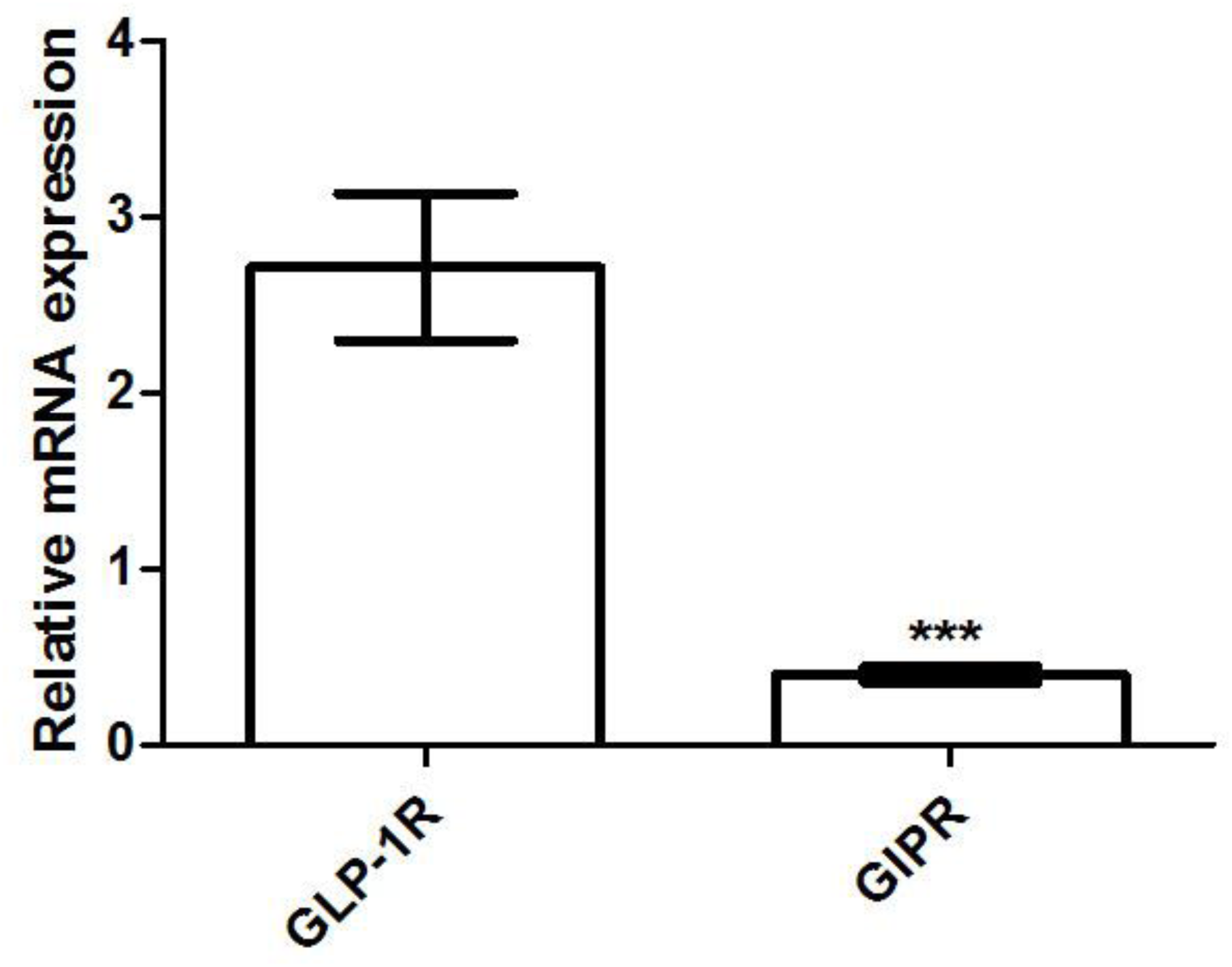
Relative mRNA expression of GLP-1R GIPR measured by the 2^-ΔΔCT^ method (Livak and Schmittgen, 2001). Data are expressed as mean ± SE; n=3; ***p<0.001 comparing GLP-1R and GIPR expression in BRIN-BD11 pancreatic beta cells.

